# Cortex-wide representational drift of different layers

**DOI:** 10.64898/2026.07.15.738614

**Authors:** Yael E. Pollak, Robert Sachdev, Matthew Larkum, Ariel Gilad

## Abstract

Representational drift, the gradual evolution of neural population codes over days, has been widely documented at the level of single neurons. However, whether drift is organized across larger cortical populations and spatial scales remains unclear. Here, we examined cortex-wide population dynamics under tightly controlled sensory input and behavior. Using widefield calcium imaging, we longitudinally recorded excitatory activity in Layers 2/3 (L2/3) or Layer 5 (L5) across 25 cortical areas while mice performed a whisker-based texture discrimination task. Sensory-evoked activity was tracked over five consecutive days during stable task performance. Population activity patterns reorganized over days, across the cortex, in both layers. This reorganization followed distinct laminar motifs: notably L5 responses exhibited a widespread, monotonic decrease in activity across most cortical areas, whereas L2/3 responses showed spatially localized and heterogeneous changes that were strongest during the sensory period. These laminar differences extended beyond the stimulus period, with L5 exhibiting more prolonged temporal engagement than L2/3. Together, these findings indicate that representational drift can unfold as a process that is coordinated across cortex with distinct laminar profiles. Thus, drift may reflect a structured feature of cortex-wide circuit dynamics.

## Introduction

Neural activity patterns are often assumed to represent stable internal representations of the external world. In sensory systems, stimuli are thought to be encoded by reliable patterns of neural activity that support consistent perception and behavior over time^1–5^. However, this assumption of stability has been challenged by studies showing that neural responses to the same stimulus can change substantially over days, a phenomenon known as representational drift (RepD;^6–11^). Importantly, the properties of this drift appear to vary across brain regions.

Earlier work has revealed several forms of representational drift. In piriform cortex^10^ and posterior parietal cortex^11^, individual neurons exhibit substantial variability across days while population-level statistics remain relatively stable. In contrast, hippocampal ensembles often reorganize more rapidly, sometimes in hours, with changes in the composition and sparsity^12,13^. In visual cortex, responses to simple stimuli such as drifting gratings remain stable across weeks, whereas responses to natural movies reorganize more strongly, indicating stimulus-dependent drift^8^. These findings have led to several distinct interpretations of drift, including the idea that drift temporally separates experiences, or reflects continual learning, generalization, or redundancy in population codes. Recent work has proposed several functional interpretations of hippocampal drift, including roles in distinguishing temporally distinct but similar experiences^6,14^, supporting continual learning^14^, enabling generalization^15,16^, and providing redundancy^17^. Conversely, drift may arise from subtle, often unmeasured, fluctuations in behaviour (e.g., running speed^18–21^) or sensory context (e.g., olfactory cues^22–24^). Supporting this latter view, studies in Egyptian fruit bats^15,25^, showed highly stereotyped flight trajectories that were associated with minimal drift in hippocampal CA1 across days^26^.

In the neocortex, most studies of representational drift have focused on individual sensory or association areas and have largely overlooked how representations change across cortical layers. Yet cortical circuits are organized in layers, with distinct connectivity and function. Superficial layers (layer 2/3) are dominated by recurrent intracortical and long-range corticocortical interactions, whereas deeper layers (layer 5) provide the bulk of the output from cortex, especially to subcortical targets^27^. Several studies suggest that drift may depend on stimulus class, cortical layer, or cell type. For example, layer-dependent differences in representational stability have been observed in visual cortex^9,28^, and receptive-field-dependent drift has been reported in barrel cortex^29^. Additional studies have described tuning instability in non-columnar somatosensory neurons^30^ and differential stability of task-related variables in parietal and retrosplenial cortex^31,32^. However, because these studies typically focus on single regions or restricted stimulus classes, it remains unclear whether RepD reflects local circuit dynamics or a coordinated process across larger cortical populations.

Computational approaches have identified several mechanisms that could give rise to representational drift. These include synaptic processes such as spike timing-dependent plasticity (STDP), synaptic turnover coupled with homeostatic normalization, stochastic synaptic-weight^33^ fluctuations driven by white^34^ or correlated^35^ noise, dropout of nodes or weights in recurrent networks^36^, and implicit regularization of population activity^13^. An alternative hypothesis posits that RepD may arise from slow fluctuations in intrinsic neuronal excitability^37^, rather than changes in synaptic connectivity. Supporting this possibility, recent work using a multisensory virtual-reality system demonstrated that hippocampal drift persisted even when sensory input and behavior were tightly controlled. Moreover, the intrinsic excitability of individual hippocampal place cells strongly predicted their future stability, with more excitable neurons exhibiting greater long-term representational stability^38^.

Together, these studies motivated us to test whether representational drift reflects local circuit dynamics or is a coordinated process that occurs across the cortex, as a whole^39–45^. To address this question, we examined cortex-wide population activity while head fixed mice performed a go-no go whisker-based texture discrimination task with well-constrained behavioral output. Using widefield calcium imaging, we longitudinally recorded excitatory activity across 25 cortical areas in either layer 2/3 (L2/3) or layer 5 (L5) over five consecutive days. This approach allowed us to test whether representational drift emerges at the population level, whether it differs across cortical layers, and to what extent behavioral and sensory variables contribute to these dynamics.

## Results

### Behavioral dynamics are stable in L2/3 and L5 transgenic mice

To quantify cortex-wide laminar representational drift independent of behavioral parameters, we recorded transgenic mice expressing GCaMP6f in L2/3 (Rasgrf-cre) or L5 (RBP4-cre) excitatory neurons across the cortex, as mice performed a whisker-based go/no-go texture discrimination task for 5 consecutive days. In short, water-restricted mice were presented with two types of sandpaper presented to the whiskers and were required to associate one texture (go) with a reward (Fig. 1a). In ‘Hit’ trials, mice were rewarded for correctly licking in response to the go texture. Incorrect licking in response to the no-go texture (‘False Alarm’, FA) resulted in a white-noise punishment, while withholding licking for either texture (‘Miss’ and ‘Correct Rejection’, CR) was neither rewarded nor punished (Fig. 1c). The trial was initiated with an auditory cue, which signaled the approach of a texture within whisker-touch for a duration of 2 s. Then, the texture moved out of position and a 2 s report window began. The trial structure remained constant across trials.

**Fig. 1.**
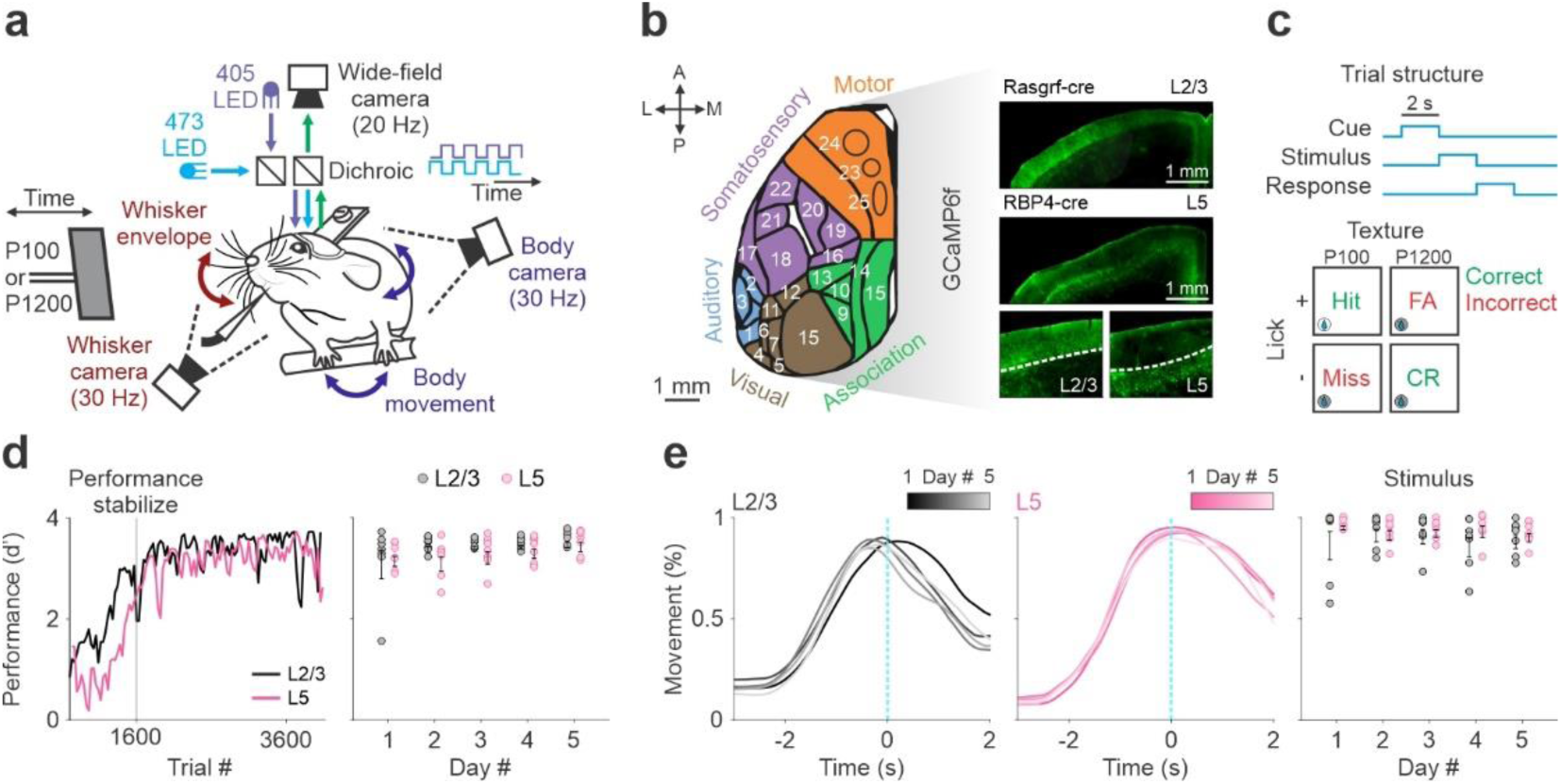
Mice groups of both L2/3 and L5 show similar behavioral profiles across days. **a** Schematic of Behavioral and imaging setup. **b** *Left:* Top view of the dorsal cortex and 25 regions of interest 1. Temporal association (TEa) 2. Auditory dorsal (AUDd) 3. Auditory primary (AUDp) 4. Post rhinal (VISpor) 5. Posterior lateral (VISpl) 6. Lateral intermediate (VISli) 7. Lateral medial (VISl) 8. Primary visual (VISp) 9. Posterior medial (VISpm) 10. Anterior medial (VISam) 11. Anterior lateral (VISal) 12. Rostro lateral (VISrl) 13. Anterior (VISa) 14. Restrosplenial angular (RSPagl) 15. Restrosplenial dorsal (RSPd) 16. Primary trunk (SSp-tr) 17. Secondary sensory (SSs) 18. Barrel cortex (SSp-bfd) 19. Primary hindlimb (SSp-ll) 20. Primary forelimb (SSp-ul) 21. Primary nose (SSp-n) 22. Primary mouth (SSp-m) 23. Whisker primary motor (MOp) 24. Anterior lateral motor (ALM) 25. Secondary motor (MOs). Regions are grouped into five divisions: Auditory (blue), Visual (brown), Association (green), Somatosensory (purple), and Motor (orange). *Right:* Enlargement of a coronal slice displaying layer-specific (L2/3 or L5) green fluorescence. **c** Trial structure and possible trial outcomes. **d** *Left:* Performance (d′) as a function of trial number averaged across mice of L2/3 (black; n = 7) and L5 (pink; n = 7). Black vertical line indicates the beginning of 5 consecutives recording days for analysis. *Right:* performance (d’) averaged across days and mice in L2/3 (black) and L5 (pink). Error bars depict mean ± s.e.m across mice. Dots depict individual mice. **e** *Left:* Movement probability (%) along the trial averaged across days (color scale) and L2/3 (black) and L5 (pink) mice during Stimulus. Vertical cyan dashed lines depict the texture stop. *Right:* Movement probability (%) averaged across days and mice in L2/3 (black) and L5 (pink). Error bars depict mean ± s.e.m across mice. Dots depict individual mice.

Once mice had learned to discriminate between textures (2–3 days after reaching a d′ of 1.5) and consistently maintained a high success rate (d′ between 2–4), we used wide-field calcium imaging to measure cortex-wide neuronal population dynamics in either L2/3 or L5 (Fig. 1b right), together with concurrent video monitoring of whisking and body movements (Methods). We imaged seven mice per layer (L2/3 or L5) over 5 consecutive days, yielding 1358-2367 trials per mouse. The Rasgrf2 line primarily labels excitatory neurons in L2/3, largely overlapping with Intratelencephalic (IT) populations. The Rbp4 line primarily labels pyramidal tract (PT) neurons in L5B but also includes a subpopulation of IT neurons in L5A. For simplicity, we refer to these lines as L2/3 and L5. Cortex-wide dynamics were aligned across days and registered onto the 2D top-view of the Allen reference atlas. Twenty-five cortical areas were defined and grouped into five general categories: Motor, Somatosensory, Association, Visual and Auditory (Fig. 1b left; Methods).

Performance levels did not differ significantly across the five days in either L2/3 or L5 mice, nor between layers (five consecutive days; ∼400 trials/day; Fig. 1d; Supplementary Fig. 1a for individual mice; n = 7 mice, two-way ANOVA; Layers *F*(14,5) = 2.21, *P* = 0.14; Days *F* = 2.24, *P* = 0.07; Layers:Days *F* = 0.64, *P* = 0.64). We also measured the mice’s body movements to determine whether their activity patterns changed over days. For each trial, we extracted forelimb and back movements from the body camera and calculated a binary movement vector by choosing a fixed threshold (Methods). Body movement, initiated as the texture approached the whiskers, did not differ significantly across the five days in either L2/3 or L5 mice, nor between layers. (during early stimulation period, i.e. Stimulus; averaged 0.4 s before and 0.1 s after texture stop; Fig. 1e; Supplementary Fig. 1b for individual mice; n = 7 mice, two-way ANOVA; Layers *F*(14,5) = 3.29, *P* = 0.08; Days *F* = 0.2, *P* = 0.94; Layers:Days *F* = 0.6, *P* = 0.66). Frame-by-frame analysis for body movement revealed similar results throughout the trial, with mostly no significant differences between days or layers (Supplementary Fig. 1b for individual mice; n = 7 mice). In summary, L2/3 and L5 mice exhibited similar and stable performance and body movements across five consecutive days, enabling investigation of representational drift in both layers under a relatively stable paradigm.

### Cortex-wide representational drift in L2/3 and L5

We analyzed the cortical activity of L2/3 and L5 as revealed by wide-field calcium imaging during Hit trials and systematically studied the characteristics of representational drift using similar parameters from previous studied^10,46^. First, we focused on the cortex wide activity during early stimulation period (i.e. Stimulus; 0.4 s before and 0.1 s after texture stop) across 5 days. Figure 2a shows example activation maps (ΔF/F) during Stimulus for L2/3 and L5, both displaying activity in SSp-bfd that is typical for this period. The mean response of L2/3 and L5 mice (n = 7) across all 25 cortical areas over consecutive recording days (n = 5) revealed varying response levels which differ across areas and layers (Fig. 2b). In general, L5 exhibited a consistent decrease in stimulus driven activity in the early epoch, across days in most cortical areas, whereas L2/3 showed more uneven inter-day fluctuations across the cortex.

**Fig. 2.**
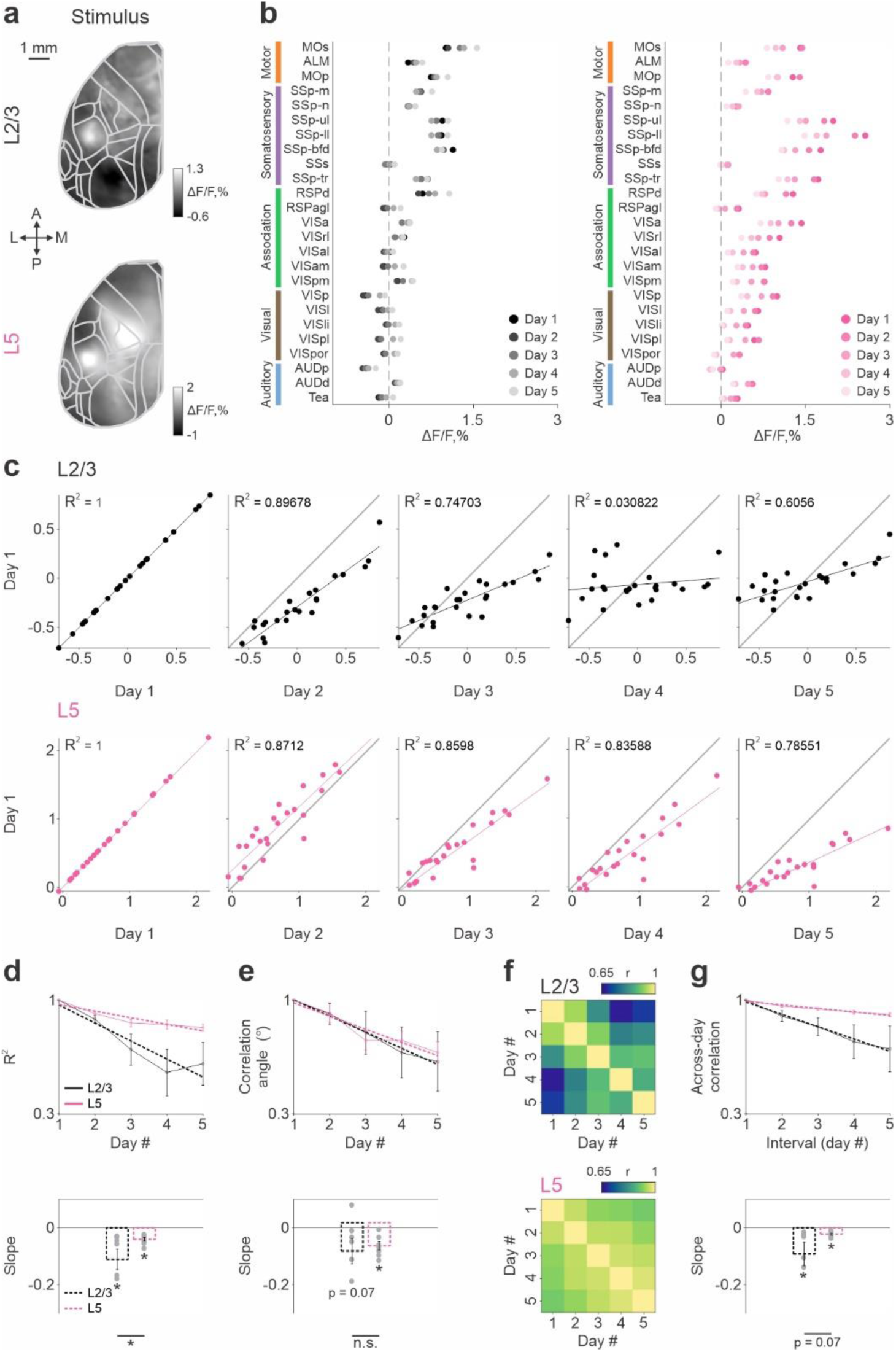
Experimental measurements reveal laminar cortex-wide representational drift during early stimulation. **a** Example activation maps from one mouse of L2/3 (top) and L5 (bottom) averaged during early stimulation (Stimulus; 0.4 s before and 0.1 s after texture stop). Color scale bar indicates min/max of percent ΔF/F. Overlay of areas in gray for all maps. Scale bar is 1 mm. **b** Mean activation of all 25 cortical areas in 5 consecutives recording days (dark to light) of L2/3 (black; n = 7) and L5 (pink; n = 7) mice during early stimulation. Areas are further divided into Motor (orange), Somatosensory (purple), Association (green), Visual (brown) and Auditory (blue) areas. **c** Scatter plot between the response in 25 cortical areas (during stimulus period) in day 1 (y-axis) and the responses in each of the five consecutive days (x-axis). Example L2/3 (black; top) and L5 (pink; bottom) mouse. R^2^ is reported for each scatter. **d** *Top:* Evolution of average R^2^ across 5 test days in L2/3 (black) and L5 (pink) mice. Dashed lines indicate slope. Error bars depict mean ± s.e.m across mice (n = 7). *Bottom:* Averaged slope across L2/3 (black) and L5 (pink) mice. Error bars depict mean ± s.e.m across mice (n = 7). Dots depict individual mice. \**p* < 0.05, signed-rank test within layer, rank-sum test between layers. **e** *Top:* Evolution of average corrected angle across 5 test days in L2/3 (black) and L5 (pink) mice. Error bars depict mean ± s.e.m across mice (n = 7). Dashed lines indicate slope. Bottom: Averaged slope across L2/3 (black) and L5 (pink) mice. Error bars depict mean ± s.e.m across mice (n = 7). Dots depict individual mice. \**p* < 0.05, n.s. not significant, signed-rank test within layer, rank-sum test between layers. **f** Average population vector correlations for 25 cortical representations of L2/3 (top) and L5 (bottom) across days. Color scale bar indicates min/max of Pearson’s correlation coefficient (r). **g** *Top:* Average correlation decay against time interval between test days in L2/3 (black) and L5 (pink) mice. Error bars depict mean ± s.e.m across mice (n = 7). Dashed lines indicate slope. *Bottom:* Averaged slope across L2/3 (black) and L5 (pink) mice. Error bars depict mean ± s.e.m across mice (n = 7). Dots depict individual mice. \**p* < 0.05, signed-rank test within layer, rank-sum test between layers.

To quantify RepD at the cortex-wide level, we correlated the average response during Stimulus across 25 cortical areas on day 1 with the corresponding average response on each subsequent day, shown for an example mouse separately for L2/3 and L5 (Fig. 2c; Pearson’s correlation, R²), as well as averaged across mice (Fig. 2d, top; n = 7 mice per layer). R² decreased as a function of days in both layers; that is, cortex-wide responses became progressively less similar across recording days when compared to the first day, indicating a gradual drift in cortex-wide response patterns over time. The slope of the change in R^2^ across days was significantly negative in both layers (*p* < 0.05; signed-rank test), with L2/3 exhibiting a significantly more pronounced negative slope than L5 (Fig. 2d bottom; *p* < 0.05; rank-sum test). In addition, the angle between the actual linear fit (comparing each day to day 1) and the perfect fit (the diagonal in Fig. 2c) decreased progressively across days, reaching significance in L5 and, to a lesser extent, in L2/3. (Fig. 2e; L5 *p* < 0.05, L2/3 *p* = 0.07; signed-rank test) with no significant difference between the two layers (*p* > 0.05; rank-sum test). Finally, the full inter-day correlation matrix (r; computed between the response vectors of 25 cortical areas across pairs of days) was calculated for L2/3 and L5, revealing high r values along the diagonal (i.e., small day intervals) and progressively lower r values with increasing distance from the diagonal (i.e., large day intervals; Fig. 2f). A similar pattern was observed across eight consecutive recording days (Supplementary Fig. 2). Pairwise correlation slope significantly decreased as a function of the interval between days in both layers (Fig. 2g; *p* < 0.05, signed-rank test), with L2/3 showing a slightly, though not significantly, stronger trend (*p* = 0.07, rank-sum test).

Next, we examined whether RepD extended to the full temporal structure of the trial. For each time frame, we computed the similarity of cortex wide activity patterns across pairs of days, using the same approach as in Figure 2g. Specifically, for each time frame, we computed the full inter-day correlation matrix and quantified the slope of the linear fit between r values and inter-day intervals (Fig. 3a). A negative slope indicates that cortex-wide activity becomes increasingly dissimilar as a function of days. In both layers, the slope exhibited pronounced temporal dynamics that differed between layers (Fig. 3b). In L5, the slope became significantly negative beginning 1.8 s before texture stop, indicating that drift is already present just after cue onset (–2 s before texture stop). This effect persisted throughout stimulus presentation and extended into the stimulus epoch, remaining significant up to 0.7 s after texture stop (*p* < 0.05; sign-rank test; FDR-corrected; Supplementary Fig. 3a shows frame-by-frame analysis. In contrast, L2/3 showed a more temporally restricted profile, with significant negative slopes observed from 1.2 s before texture stop up to texture stop (time 0), indicating that effects were predominantly confined to the stimulus period. A direct comparison between layers did not reveal significant differences in slope at any time point (*p* > 0.05; rank-sum test; FDR-corrected; Supplementary Fig. 3b). Together, these results indicate that cortex-wide representational drift is not solely a stimulus-locked phenomenon but extends across the trial, with a more prolonged engagement in L5.

**Fig. 3.**
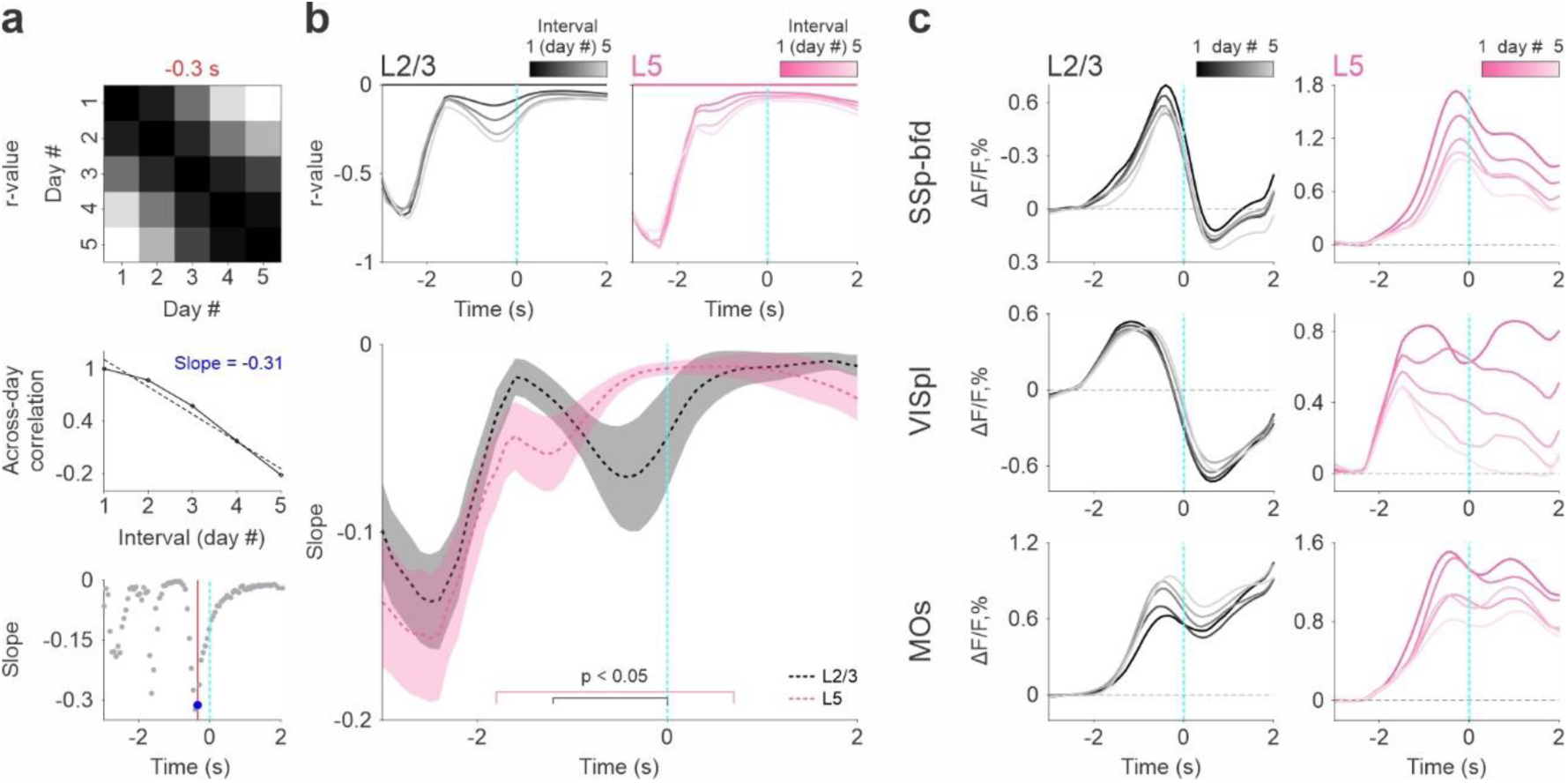
Laminar cortex-wide representational drift across the trial. **a** Schematic illustration for calculating cortex-wide slope of correlation decay against time interval between test days for each time frame. **b** *Top:* Correlation decay curve against time interval between test days (dark to light) averaged across L2/3 (black; n = 7) and L5 (pink; n = 7) mice. *Bottom:* Slope of correlation decay curve against time interval between test days averaged across L2/3 (black) and L5 (pink) mice. Error bars depict mean ± s.e.m across mice (n = 7). Vertical cyan dashed lines depict the texture stop. Bars indicate *p* < 0.05, signed-rank test; FDR-corrected for L2/3 (black) and L5 (pink). **c** response curve ΔF/F of three selected areas: SSp-bfd, VISpl and MOs (from top to bottom) averaged across 5 days (color scale) and L2/3 (black; n = 7) or L5 (pink; n = 7) mice. Vertical cyan dashed lines depict the texture stop. Horizontal black dashed line depicts 0 response.

Finally, to relate the population-level measures to area-specific dynamics, we examined the mean temporal response profiles in representative cortical regions, including barrel cortex (SSp-bfd), posterior lateral cortex (VISpl), and secondary motor cortex (MOs), all of which have been shown to be involved in this task^42^. Consistent with the analysis in Fig. 3b, L5 responses across all three areas showed a gradual decrease over time but with a varying extent. In contrast, L2/3 responses were more heterogeneous: SSp-bfd exhibited an overall increase, VISpl displayed a biphasic profile with an initial decrease followed by a rise during stimulation, and MOs showed a general increase. Together, these findings suggest that the extended representational drift observed at the population level arises from distinct and non-uniform changes across L2/3, whereas L5 exhibit more coherent modulation across areas.

### Representational drift is local in L2/3 and global in L5

To examine how representational drift is distributed across cortical areas, we analyzed layer-specific activity maps (ΔF/F) during the Stimulus period over five consecutive days (Fig. 4a). Comparison of day-to-day activity maps revealed a consistent decrease in L5 activity over time, whereas activity in L2/3 changed in a more heterogeneous manner. Extended activity maps spanning seven consecutive recording days showed similar results, further demonstrating the continuous progression of the changes described above (Supplementary Fig. 4a). To quantify these effects, we compared mean responses between the first 300 (early) and last 300 (late) trials for each layer (Fig. 4b). In L2/3, activity increased significantly in association and motor areas, including VISpl and MOs (*p* > 0.05; signed-rank test; FDR-corrected), whereas somatosensory areas such as SSp-bfd remained largely unchanged (*p* > 0.05; signed-rank test; FDR-corrected). In contrast, L5 showed widespread suppression, with significant decreases across most somatosensory, association and visual areas (*p* < 0.05; signed-rank test; FDR-corrected), with only a few exceptions. Together, these results indicate that activity changes in L2/3 are spatially heterogeneous, whereas L5 exhibits a broad and more uniform decrease in activity.

**Fig. 4.**
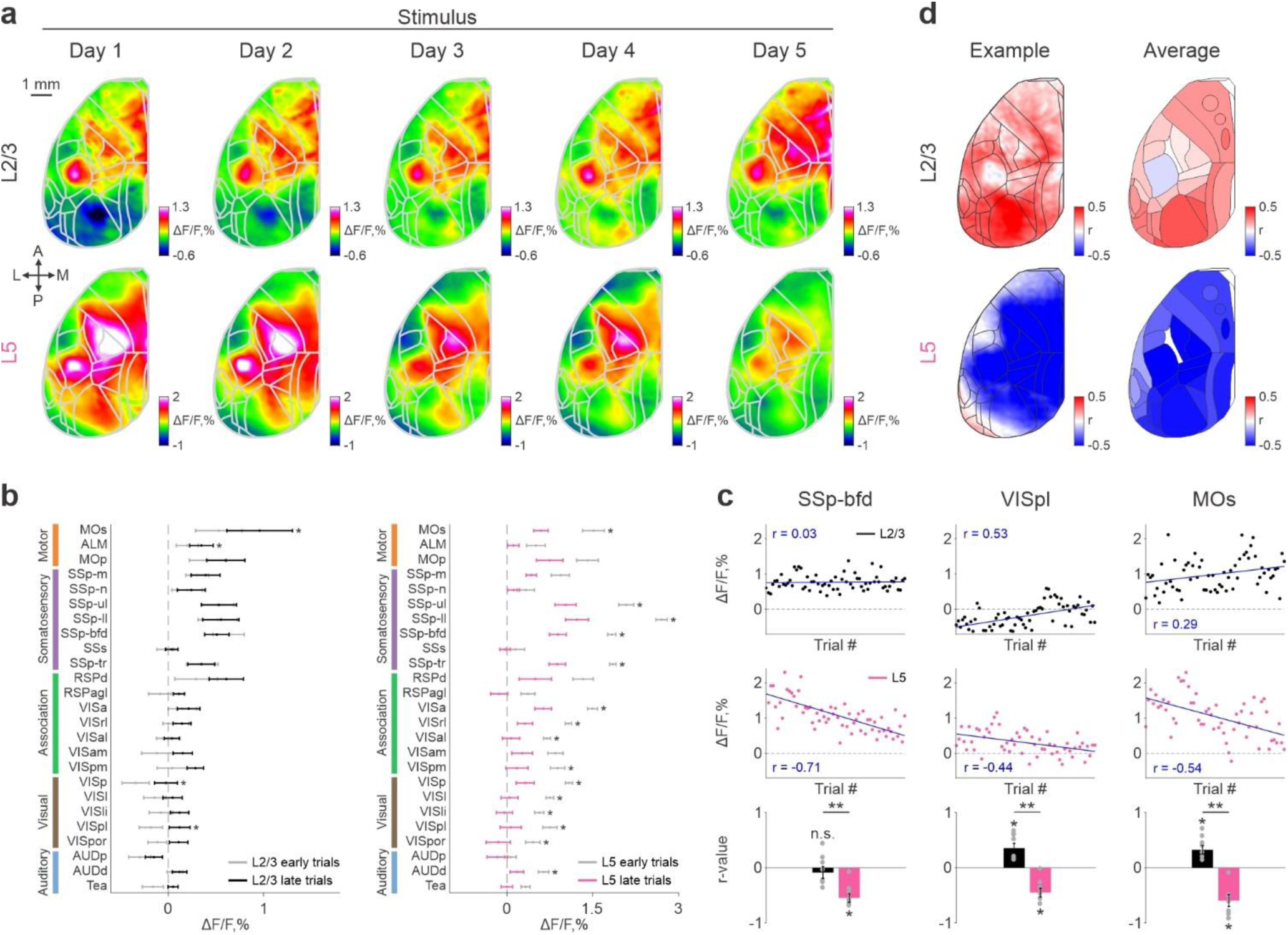
Laminar representational drift in relationship to trial number dissociate during early stimulation. **a** Example activation maps from one mouse of L2/3 (top) and L5 (bottom) averaged during early stimulation (Stimulus) and across 5 consecutive days (left to right). Color scale bar indicates min/max of percent ΔF/F. Overlay of areas in gray for all maps. Scale bar is 1 mm. **b** Mean activation of all 25 cortical areas in early (gray) and late (color) trials of L2/3 (black) and L5 (pink) mice during early stimulation. Error bars depict mean ± s.e.m across mice (n = 7). \**p* < 0.05, Signed-rank test. FDR-corrected. **c** *Top:* Mean responses across trials (binned every 30 trials) for three selected areas: SSp-bfd, VISpl and MOs (from left to right) averaged during early stimulation in example L2/3 (black) and L5 (pink) mouse. *Bottom:* Pearson’s correlation coefficient (r) for each area during Stimulus averaged across L2/3 (black) and L5 (pink) mice. Error bars are ± s.e.m. across mice (n = 7). Dots depict individual mice. \**p* < 0.05, ** *p* < 0.005, n.s. not significant, signed-rank test within layers, rank-sum test across layers, FDR-corrected. **d** *Left:* Pearson’s correlation coefficient (r) for each pixel of example L2/3 (black) and L5 (pink) mouse (same to those presented in a) during early stimulation. Color scale bar indicates min/max of Pearson’s correlation coefficient (r). *Right:* similar to figure on the left this time averaged across all 25 cortical areas and L2/3 (black) and L5 (pink) mice. Color scale bar indicates min/max of Pearson’s correlation coefficient (r).

To quantify RepD within each cortical area, we computed Pearson’s correlation coefficient of response activity as a function of time (r; response vs. trial bins) during the Stimulus period (Fig. 4c). Positive r values indicate increasing activity over time, negative values indicate decreasing activity, and values near zero indicate temporal stability. Focusing on the three representative areas (SSp-bfd, VISpl, and MOs) we observed a consistent pattern in L5: all three areas exhibited significant negative r values (Fig. 4c; *p* < 0.05; signed-rank test), indicating a progressive decrease in activity over trials. In contrast, L2/3 exhibited heterogeneous region-specific dynamics: SSp-bfd showed no significant drift (*p* > 0.05; signed-rank test), whereas VISpl and MOs displayed significant positive r values (*p* < 0.05; signed-rank test), reflecting increasing activity across trials. Notably, in all three regions, r values were significantly different between layers (*p* < 0.005; rank-sum test), suggesting that temporal trends in the both layers evolved differently. We extended this analysis to the entire dorsal cortex by computing r values for each cortical area and each pixel (Fig. 4d). This analysis revealed striking, layer-specific spatial patterns, L2/3 exhibited predominantly positive r values across both frontal and posterior regions, whereas L5 exhibited negative r values across nearly all cortical areas (Fig. 4d; see also Supplementary Fig. 4b). Together, these results indicate that representational drift in L2/3 and L5 during the Stimulus epoch differs markedly in its spatial organization across cortical areas. In L2/3, drift is region-specific and spatially heterogeneous, reflecting localized changes in activity. In contrast, L5 exhibits a more global and spatially uniform drift, characterized by a widespread reduction in activity across areas.

Next, we applied this analysis beyond the Stimulus period to encompass the full temporal structure of the trial (Fig. 5). For each time frame, we computed Pearson’s correlation coefficient of response activity as a function of time (r; response vs. trial bins), yielding a temporal vector of drift that captures the magnitude and direction of changes in activity across trials within the temporal trial structure (Fig. 5a). Focusing on the three representative areas (SSp-bfd, VISpl, and MOs), we first visualized activity as two-dimensional maps of response amplitude (x-axis) versus trial bins (y-axis) over time (trial bins), revealing complex but structured changes that evolved across both short (seconds) and long (days) time scales (Fig. 5b, top). The r value (described above and in Fig. 5a) yielded a time resolved measure of RepD that captures both the direction and the magnitude of changes over days (Fig. 5b, bottom). In L5, all three areas exhibited negative r values shortly after cue onset and persisting beyond texture stop, indicating a sustained decrease in activity across trials throughout the trial time course. In contrast, L2/3 showed more heterogeneous temporal profiles: VISpl and MOs displayed positive r values primarily around the stimulus period, whereas SSp-bfd exhibited predominantly near-zero r values. A comprehensive analysis across all 25 cortical areas confirmed these patterns across the cortex (Supplementary Fig. 5). Together, these results demonstrate that cortex-wide response drift across trials unfolds in a complex, spatiotemporal manner that differs across cortical areas and cortical layers.

**Fig. 5.**
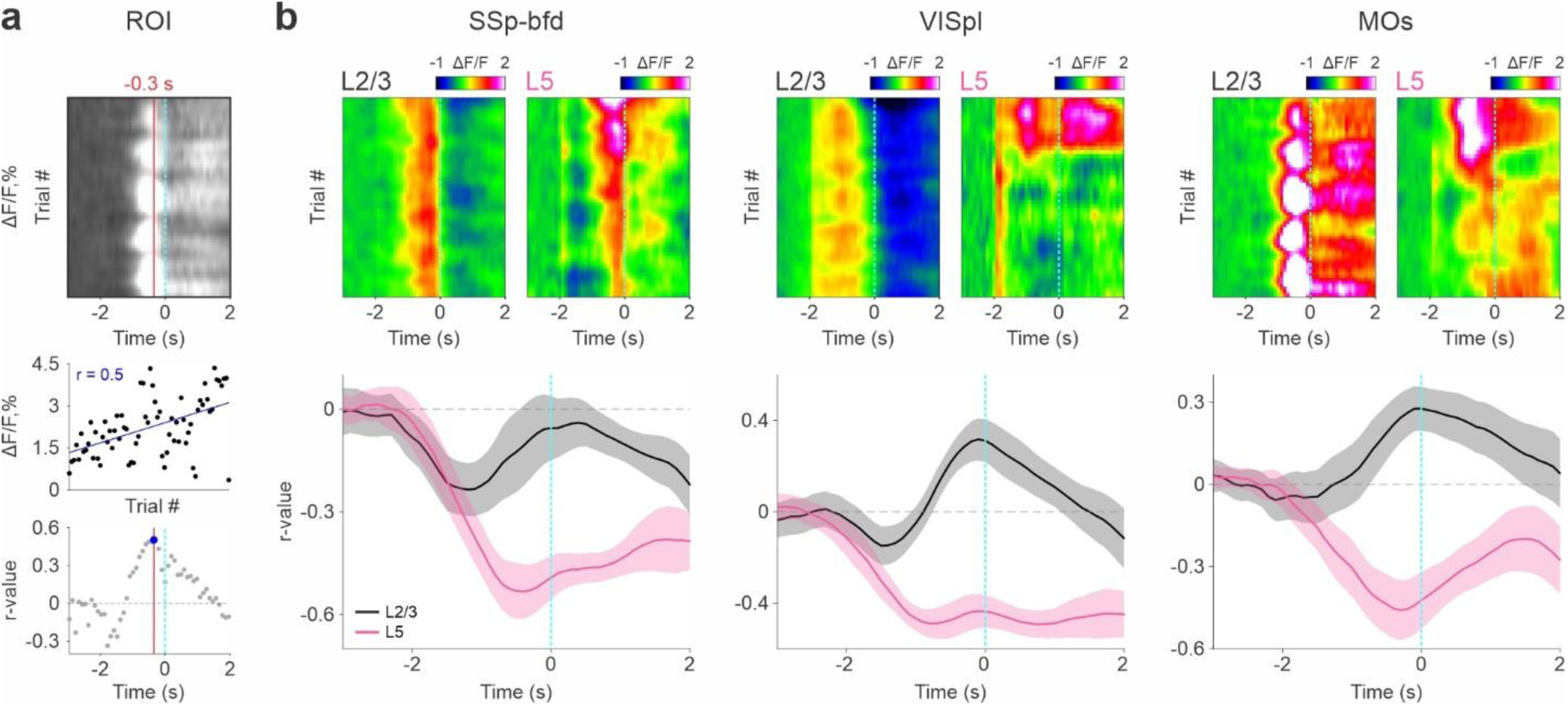
Laminar representational drift temporal relation to trial number reveals diverged dynamics. **a** Schematic illustration for calculating the Pearson’s correlation coefficient (r) across trials for each time frame for a specific ROI. b Top: Example of a 2D heat map of response (ΔF/F) from one L2/3 (left) and L5 (right) mouse (trial structure along x-axis; trial scale along y-axis) for three areas of interest: SSp-bfd, VISpl and MOs (from left to right). Data is binned every 30 trials. Cyan vertical dashed line indicates texture stop. Color scale bar indicates min/max of percent ΔF/F. Bottom: Pearson’s correlation coefficient (r) curve averaged across L2/3 (black) and L5 (pink) mice. Error bars are ± s.e.m. across mice (n = 7). Vertical cyan dashed lines depict the texture stop.

Given the high dimensionality of spatial and temporal changes across layers, we examined whether a lower-dimensional structure could capture representational drift across layers. To this end, we applied t-distributed Stochastic Neighbor Embedding (t-SNE) to the temporal r profiles across 25 cortical areas within each layer (i.e., 90 × 25 matrix: time frames × cortical areas presented in Supplementary Fig. 6a) and generated a low-dimensional representation indicating similarities and differences in representational drift across time, cortical areas and layers. Similar temporal profiles were generated with CR trials for each layer, both showing almost identical patterns to those observed in the Hit trials temporal profiles used in the current analysis (Supplementary Fig. 6b). The resulting t-SNE embedding revealed patterns that were not explained by the laminar profile (Fig. 6a; Methods). Color-coding clusters by cortical region (Motor, Somatosensory, Association, Visual and Auditory) showed that each cluster comprised a heterogeneous mix of cortical areas (Fig. 6a). We subsequently applied k-means clustering with the elbow method to identify three clusters within this low-dimensional space (Fig. 6a; Methods). The average drift (r) was quantified within each cluster across mice (n = 7) for both layers (Fig. 6b). Cluster 1 showed no significant drift in either layer (*p* > 0.05; signed-rank test) and no significance difference between layers (*p* > 0.05; rank-sum test). Cluster 2 exhibited significantly negative drift in L5 (*p* < 0.05; signed-rank test) and a non-significant negative trend in L2/3 (*p* = 0.07; signed-rank test). In contrast, cluster 3 revealed a clear divergence between layers: L2/3 showed significantly positive r values, whereas L5 showed significantly negative values (*p* < 0.05; signed-rank test). with significant differences between layers in cluster 2 and cluster 3 (*p* < 0.01; rank-sum test).

**Fig. 6.**
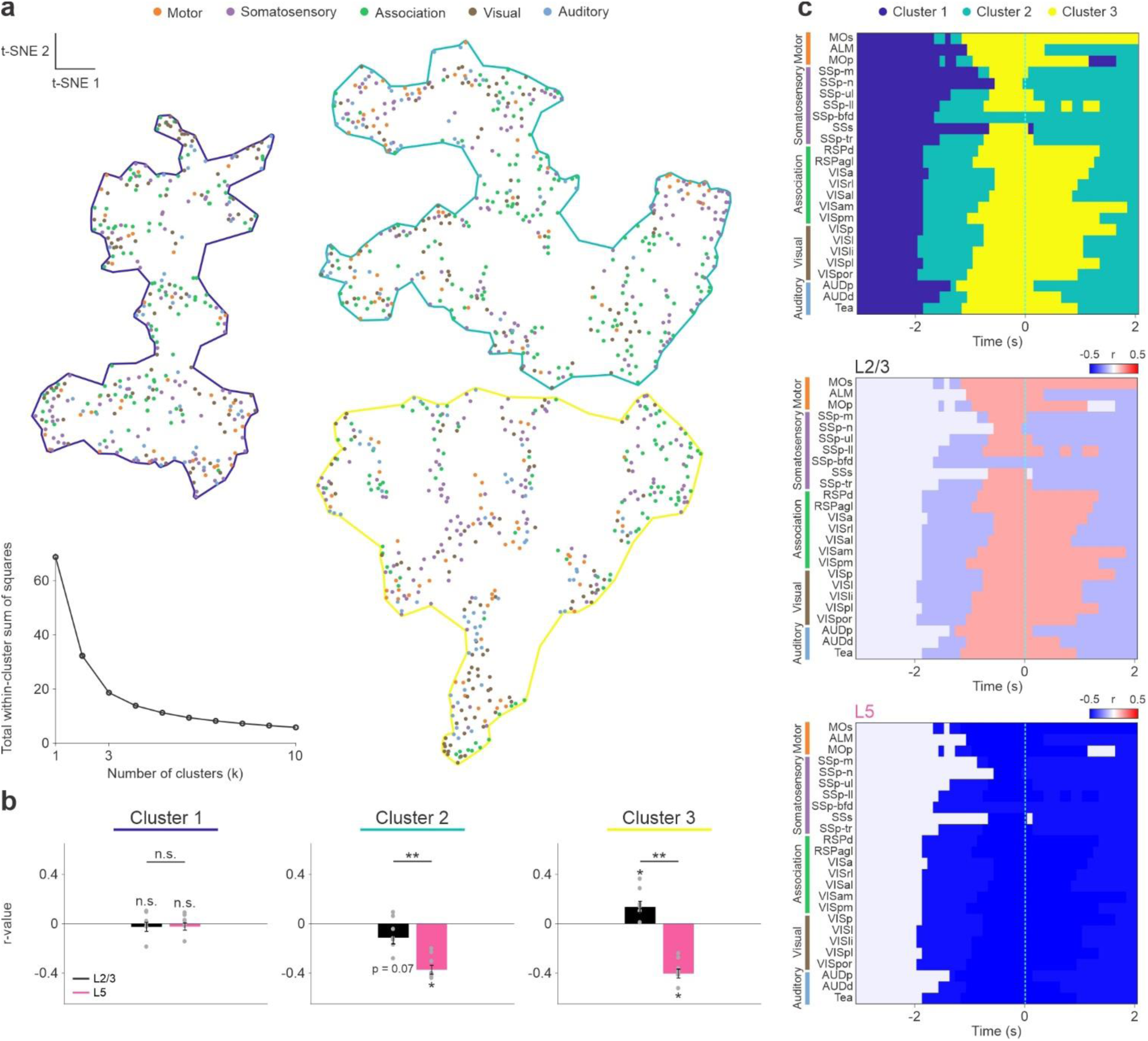
Dimensionality reduction reveals cross laminar representational drift patterns. **a** Two-dimensional t-SNE of layers using 25 cortical areas as Pearson’s correlation coefficient (r) curve averaged across L2/3 and L5 mice (25 x 90). Inset depicts an elbow plot displaying the total within-cluster sum of squares as a function of number of clusters, indicating an optimal number of clusters (k = 3). b Pearson’s correlation coefficient (r) averaged across each and L2/3 (black) and L5 (pink) mice. Error bars are ± s.e.m. across mice (n = 7). Dots depict individual mice. \**p* < 0.05, ** *p* < 0.005, n.s. not significant, signed-rank test within layers, rank-sum test across layers. c *Top:* 25 cortical areas during each time frames across the trial color coded for different clusters (k = 3). *Bottom:* 25 cortical areas during each time frames across the trial color coded for the averaged Pearson’s correlation coefficient (r) of L2/3 (top) and L5 (bottom) of different clusters (k = 3) as presented in d. Color scale bar indicates min/max of Pearson’s correlation coefficient (r). Vertical cyan dashed lines depict the texture stop.

Next, we examined the temporal and cortical distribution of each cluster by assigning each time frame in each cortical area its cluster identity (Fig. 6c; i.e., a 2D plot of time by cortical areas color coded for the three clusters). This analysis revealed a distinct distribution for each cluster in spatiotemporal space. Cluster 1 dominated the baseline period across all cortical areas and persisted until cue onset (2 s before texture stop). Cluster 2 first emerged in association and visual areas and persisted through the cue period until shortly before texture stop (−1 s). Cluster 3 emerged during the stimulus period, excluding SSp-bfd, and remained active in motor, association and visual areas for ∼ 1 s after texture stop. Following stimulus offset, most cortical areas reverted to cluster 2 with the exception of MOs. Somatosensory regions also transitioned to cluster 2 more rapidly than other cortical areas. Next, we color coded the 2D matrix based on the average r value for each cluster (Fig. 6b) for each layer separately which resulted in two distinct layer-specific drift patterns (Fig. 6c bottom). L5 showed globally negative r values (combined for both cluster 2 and 3) across all 25 cortical areas, consistent throughout the trial, beginning at cue onset in association and visual areas, followed shortly by auditory and motor areas, and lastly somatosensory areas. In contrast, L2/3 displayed a dynamic progression, transitioning from negative r values, particularly in association areas after cue onset, to positive r values during early stimulus (excluding SSp-bfd), before returning to negative values post-stimulus (excluding MOs). These findings further support the layer-specific temporal divergence observed in prior analyses. Together, they reveal distinct representational drift dynamics in each layer: L5 exhibits a global, sustained suppressive drift characterized by response diminution over days, whereas L2/3 fluctuates between negative and positive drift in a stimulus-dependent manner. These layer-specific changes in activity with time (i.e., representational drift), raise the question whether the observed changes reflect behavioral variability or arise from processes that evolve independently of behavior.

### Cortex-wide representational drift is driven by time, not behavior

To determine whether changes in behavior parameters could explain the observed neuronal drift, we applied a multivariate linear model with ridge regularization. This model incorporates ten behavioral and task-related predictors, including trial number, performance (d′), movement probability, lick and whisk movement, success, choice outcome (Hit, Miss, CR, FA), stimulus type (P1200 or P100), previous success and previous choice history (Supplementary Fig. 7a; see Methods^41^). The model was fit separately for each cortical layer, brain area, time window, and individual mouse. The resulting coefficients (β) quantify the strength and direction of the relationship between a given predictor and neuronal activity, with non-zero β-values indicating a significant contribution of that predictor. We first examined the β coefficients corresponding to the trial number predictor across time and cortical areas (25 areas × 90 frames) for each layer (Fig. 7a; n = 7 mice for each layer). These patterns closely mirrored the neuronal drift patterns quantified by r values shown in Figure 6c. In the three representative areas (SSp-bfd, VISpl and MOs), the temporal evolution of β coefficients recapitulated the respective drift profiles from Figure 6b, with the consistently negative coefficients in L5 and positive coefficients in L2/3 around the stimulus period. When averaged over the Stimulus period and visualized for the entire dorsal cortex, L5 exhibited widespread negative β values, whereas L2/3 showed significantly positive β values in a subset of cortical areas (Fig. 7b; see also Supplementary Fig. 7c). Together, these results demonstrate that that trial number predictor, which serves as a proxy for time, robustly predicts neuronal responses in both layers, with a stronger effect and more uniform effect observed in L5.

**Fig. 7.**
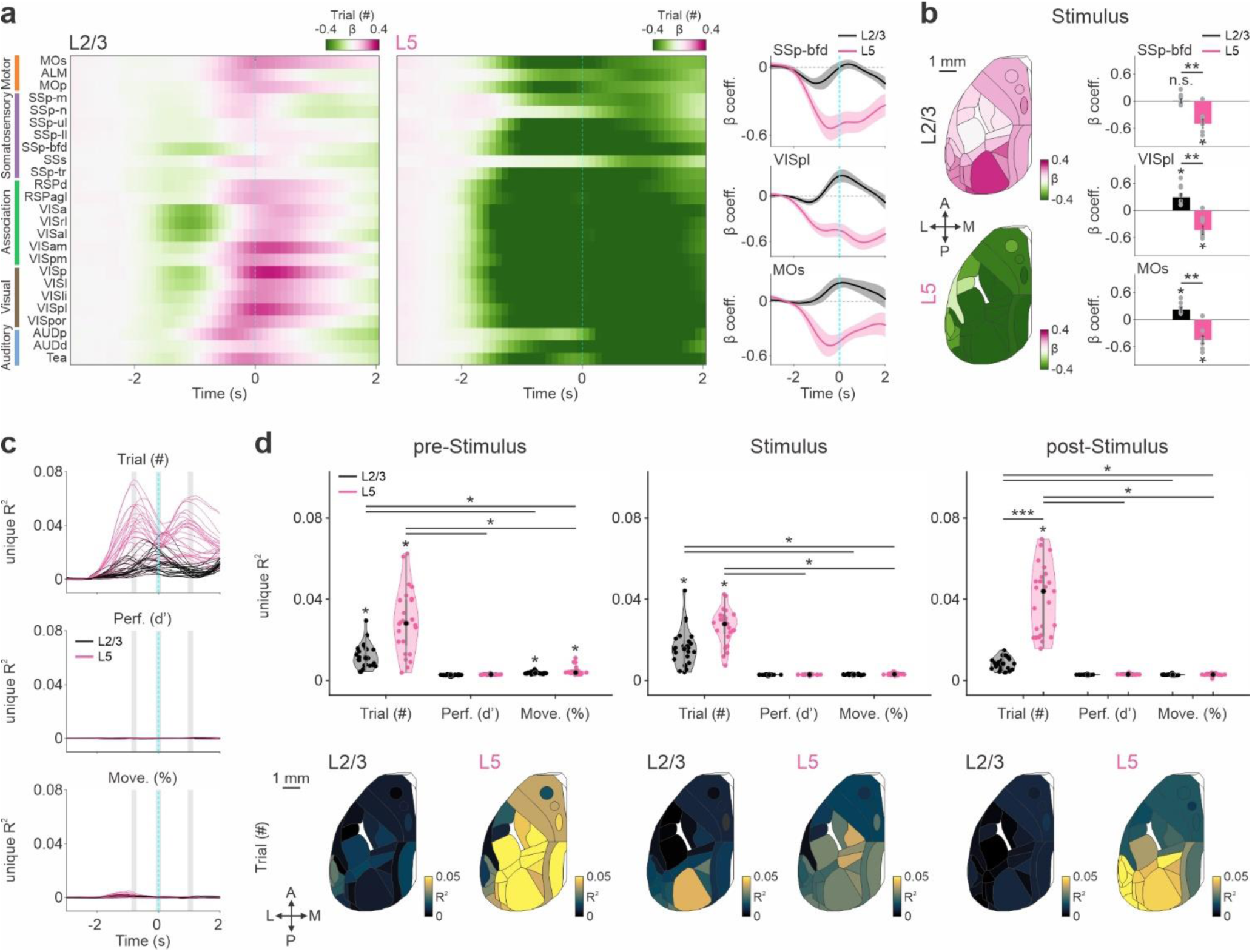
Laminar representational is largely explained by time. **a** *Left:* Coefficient value (β) averaged for the trial number predictor (trial (#)) as cortical activity patterns across time averaged across 25 cortical areas and L2/3 (left) and L5 (right) mice. Color scale bar indicates min/max of Coefficient value (β). Vertical cyan dashed lines depict the texture stop. *Right:* Coefficient value (β) curve of three selected areas: SSp-bfd, VISpl and MOs (from top to bottom) averaged across L2/3 (black) and L5 (pink) mice. Error bars are ± s.e.m. across mice (n = 7). Vertical cyan dashed lines depict the texture stop. Horizontal black dashed line depicts r value = 0. **b** *Left*: Coefficient value (β) averaged across all 25 cortical areas and L2/3 (black) and L5 (pink) mice during early stimulation (Stimulus). Color scale bar indicates min/max of Coefficient value (β). *Right:* Coefficient value (β) of three selected areas: SSp-bfd, VISpl and MOs (from top to bottom) averaged across L2/3 (black) and L5 (pink) mice during early stimulation (Stimulus). Error bars are ± s.e.m. across mice (n = 7). Dots depict individual mice. \**p* < 0.05, ** *p* < 0.005, n.s. not significant, signed-rank test within layers, rank-sum test across layers, FDR-corrected **c** unique R^2^ curve of three predictors: performance (Perf. d’), movement probability (Move. %) and trial number (Trial. #), from top to bottom, averaged across 25 cortical areas and L2/3 (black) and L5 (pink) mice. Gray bars indicate selected temporal periods presented in d. Vertical cyan dashed lines depict the texture stop. **d** *Top:* Violin plot of unique R^2^ averaged across three selected temporal periods: pre-Stimulus, Stimulus and post-Stimulus (from left to right) for the trial number (Trial. #), performance (Perf. d’) and movement probability (Move. %) predictors for L2/3 (black) and L5 (pink) mice. Dots depict 25 different brain areas averaged across mice (n = 7). \**p* < 0.05, \*\*\**p* < 0.0005, signed-rank test within layer, rank-sum test across layers. Bonferroni corrected. *Bottom:* cortical maps of unique R^2^ for the trial number predictor averaged across three selected temporal periods: pre-Stimulus, Stimulus and post-Stimulus (from left to right) and L2/3 (left) and L5 (right) mice. Color scale bar indicates min/max of unique R^2^.

Next, we examined the unique contribution of specific behavioral parameters, focusing on trial number, performance (d′) and movement probability^42,47^. To isolate the unique contribution of each predictor, we computed the unique R² by subtracting the R² of a model containing a shuffled version of a given predictor from the R² of the full model (see Methods). Unique R² values for performance and movement remained low across all cortical regions and time points in both layers (Fig. 7c). In contrast, the trial number predictor displayed high unique R^2^ for several cortical areas in both layers. Notably, there was a distinct temporal structure for the two layers: two peaks emerged over the course of the trial in L5 whereas in L2/3 there was a single peak around the Stimulus period (Fig. 7c, bottom). To quantify these dynamics, we averaged unique R² values across all 25 cortical areas and mice within each layer for three epochs: pre-Stimulus (–0.8 to –0.7 s before texture stop), Stimulus (–0.1 s up to texture stop), and post-Stimulus (0.9 to 1 s after texture stop; Fig. 7d). Across these epochs, performance and movement predictors did not differ from zero (*p* > 0.05; signed-rank test; Bonferroni corrected), with the exception of a significant effect of movement in both layers during the pre-Stimulus epoch (*p* < 0.05; signed-rank test; Bonferroni corrected). In contrast, the trial number predictor exhibited significant unique contributions in both layers across all epochs (*p* < 0.05; signed-rank test; Bonferroni corrected). Layer specific differences emerged in the post-Stimulus epoch, with L5 exhibiting significantly higher unique R² values than L2/3 (*p* < 0.001; rank-sum test; Bonferroni corrected). unique R² values for individual predictors for each time epoch are provided in Supplementary Fig. 7b.

Mapping these values onto the cortical surface revealed a dynamic layer specific sequence of engagement (Fig. 7d, bottom). During the pre-Stimulus epoch, L5 exhibited widespread contributions across the cortex. In the Stimulus period, L2/3 contributions became more prominent, particularly in posterior regions, along with a widespread and global reduction in L5 contributions. In the post-Stimulus epoch, L2/3 contributions declined, while L5 impact increased again, particularly in posterior areas. Averaged unique R² values across all 25 cortical areas for L2/3 and L5, along with detailed statistical comparisons, are provided in Supplementary Fig. 7d. Together, these results identify a trial progression as the dominant predictor of cortical activity across areas and layers. This effect is strongest in L5 where it unfolds with distinct temporal structure, suggesting that cortex wide dynamics are organized by internal time dependent variables rather than movement or behavioral performance variables.

It is important to note that trial number is an index of the animal’s progression through the session; its dominant contribution indicates that a large fraction of cortical activity reflects slow, time-dependent changes in activity, rather than any changes in immediate sensory input, movement, or behavioral outcome. This pattern is consistent with cortex-wide representational drift, in which population activity evolves gradually over time. The stronger and more sustained contribution of trial number in L5 further suggests that deep-layer circuits preferentially track and express these slow dynamics, whereas L2/3 activity is more tightly coupled to stimulus-related processing. Together, these results provide direct support for the view that representational drift is a dominant organizing principle of large-scale cortical dynamics (see Discussion).

Finally, we present the β coefficients for the movement predictor across time and cortical areas (25 areas × 90 frames) for each cortical layer (Supplementary Fig. 7e). Notably, prior to texture stop, positive β coefficients were observed predominantly in motor regions, including MOs and MOp, as well as in somatosensory regions such as SSp-ul and SSp-bfd. We next examined the unique R² values during the pre-Stimulus period and observed a positive trend in frontal cortical areas (Supplementary Fig. 7f). Together, these findings, consistent with previous results reported by Oz Rokach et al 2025, indicate that movement predicts neuronal responses within a temporally restricted phase and in a region-specific manner, with the strongest effects localized to frontal cortical areas traditionally associated with movement-related processing.

## Discussion

Here we demonstrate that there is a laminar dissociation in representational drift that is temporally driven, present at the population level, and independent of behavior. Over days, the Population activity patterns reorganize distinctly in both layers with L2/3 exhibiting spatially localized, heterogeneous, and stimulus-linked drift, whereas L5 displays a widespread and predominantly suppressive drift that extends beyond the sensory epoch. Importantly, our work shows that this reorganization in cortical activity is related to trial number, time in a session, and it is not related to movement or behavioral parameters.

### Representational drift emerges at the cortex-wide population level

Recent technical advances have enabled longitudinal tracking of neuronal activity over days and weeks^11,14,48^. Most long-term studies have focused on single neurons within restricted brain regions, including the hippocampus^6,7^, primary visual cortex^8,9^, and posterior parietal cortex^9,11^. These studies consistently show that neuronal tuning properties can gradually change: neurons may lose responsiveness to a stimulus while others acquire it, reflecting pronounced single-cell representational drift across contexts, layers, and cell types. Notably, however, many of these studies report that population-level statistics remain relatively stable despite substantial single-neuron variability^6–8,10,11,49,50^. Stable decoding performance across days has further supported the view that population representations preserve task-relevant information even as individual neurons drift^51^. Thus, single-cell instability and population-level stability have often been considered complementary, reconciling cellular plasticity with reliable stimulus representation and behavior.

In parallel, large-scale recording approaches have enabled tracking of cortex-wide activity at both single-cell resolution^52^ and at the mesoscale using wide-field imaging (Gilad and Helmchen 2020; Pollak et al. 2025; Nietz et al., 2023; Makino and Komiyama 2017). Many of these studies emphasize learning-related modulation across cortical areas and trial structure. However, such changes are typically interpreted in relation to performance, learning stage, or behavioral variability. Day-to-day neuronal reorganization that is invariant to behavioral parameters, i.e., representational drift per se, has not been systematically examined at the cortex-wide population level.

Here, under stable behavioral performance across five consecutive days, we show that the population vector derived from 25 cortical areas changes progressively over time. This reorganization becomes evident only when examining cortex-wide activity patterns, as local recordings alone would not capture the distributed population shift. Importantly, cortex-wide population drift does not contradict the existence of single-cell drift; rather, it reveals an additional mesoscale dimension of representational reorganization. Whether similar cortex-wide drift extends to subcortical circuits, other sensory modalities, or freely moving behavior remains an important direction for future work.

### Laminar organization of cortex-wide representational drift

Previous studies have reported representational drift in both L2/3 and L5 within individual cortical regions, but clear qualitative differences between layers have not been evident^8,9^. Here we demonstrate that although both layers exhibit cortex-wide drift, the nature of this drift differs along three key dimensions: magnitude and direction, spatial distribution, and temporal profile.

First, the dynamics of drift differ across layers. L2/3 exhibits greater heterogeneity across cortical areas, with many regions showing gradual enhancement over days, in contrast, L5 displays a broadly suppressive trend of varying magnitude. Second, the spatial extent of drift diverges across layers. In L2/3, enhancement is primarily localized to posterior association and visual areas with more limited changes in primary somatosensory cortex. In contrast, L5 suppression spans association, somatosensory, and motor areas. These broad differences in the patterns of representational drift indicate that representational drift in L2/3 is spatially localized, whereas in L5 it is broadly distributed. Third, in both L2/3 and L5, the temporal profile of drift differs within the trial structure. In L2/3, drift emerges predominantly during stimulus presentation, suggesting a close link to sensory representation. Whereas L5 drift begins just after cue onset and extends into the report period, indicating engagement beyond sensory encoding.

This broader temporal footprint suggests that L5 drift may relate to a habitual shift that less involves the cortex and shifts processing towards alternative subcortical structures such as the basal ganglia^53–56^. Together, these findings indicate that representational drift is not a uniform phenomenon across layers but instead reflects layer-specific modes of circuit reorganization operating at distinct spatial and temporal scales.

### Functional significance of population-level representational drift

At the single-cell level, it remains debated whether representational drift reflects biological noise or a functionally meaningful process^11,14,48,57,58^. At the population level, our findings suggest that representational drift represents coordinated network reorganization rather than random variability. The localized, stimulus-linked enhancement observed in L2/3 is consistent with adaptive reweighting within recurrent cortical circuits^40–42^. Superficial layers are dominated by intracortical and corticocortical connectivity, making them well positioned for experience-dependent refinement. We speculate that gradual enhancement in higher-order association areas may reflect circuit-level optimization of sensory integration. One potential mechanism could involve long-term modulation of inhibitory control, for example via somatostatin interneurons, leading to decreased inhibition of local L2/3 populations. The pronounced drift in posterior association areas such as VISpl, VISl, and VISli, regions implicated in higher-order sensory processing (Oz Rokach et al 2025; Pollak et al 2025; Gilad et al. 2018; Gallero Salas et al. 2021; Deitch, Rubin & Ziv, 2021; Schoonover et al., 2021), further supports the idea that superficial-layer drift reflects refinement of associative representations. In contrast, the relative stability of SSp-bfd suggests that lower-order sensory representations remain comparatively constrained.

By contrast, the widespread suppression observed in L5 suggests a different functional role. Deep-layer pyramidal neurons constitute the principal cortical output to subcortical and motor structures^59–63^. During expert performance, broad cortical output may initially support sensory processing, feedback, and motor planning. Over time, however, redundant or distributed output may become unnecessary. The gradual suppression of L5 responses across multiple areas may therefore reflect increasing efficiency of cortical output, minimizing redundant signaling while preserving essential motor commands. The relative stability of frontal areas in L5 may indicate preservation of motor-related signals, consistent with limited drift in motor regions^64–67^.

More broadly, the reduction of L5 activity may signal a shift in network recruitment during extended expert performance. Habitual or well-trained behaviors are thought to involve increased engagement of subcortical circuits, including the basal ganglia^68–70^. Under this view, prolonged task execution may rely less on distributed cortical output and more on subcortical pathways, with cortex maintaining a more streamlined role. We hypothesize that specific subpopulations of L5 neurons in the frontal cortex, particularly those projecting to striatum, may remain stable, whereas others progressively reduce their engagement. Importantly, representational drift in both layers emerged despite stable behavioral performance. This dissociation suggests that stable behavior does not require fixed population activity patterns. Instead, task-relevant information may be preserved within low-dimensional subspaces or distributed circuit architectures, even as the precise distribution of activity across areas shifts. Alternatively, compensation by other cortical layers, inhibitory subtypes, or subcortical circuits may buffer behavioral output against cortical reorganization.

The observation of continued drift over extended timescales indicates that representational reorganization is an ongoing process, even during stable expert performance. These findings suggest that cortical circuits remain dynamically reconfigurable rather than settling into a fixed steady state. Such flexibility may support robustness to synaptic turnover, metabolic constraints, or internal state fluctuations, while preserving functional output. Future studies should examine whether similar cortex-wide laminar drift occurs across sensory modalities, in freely moving behavior, and across additional cortical and subcortical structures. Abrupt changes in task demands may also reveal how rapidly these laminar motifs reorganize. In summary, our findings indicate that representational drift extends to the cortex-wide population level and follows layer-specific organizational principles. Rather than reflecting noise, representational drift may represent an intrinsic mode of circuit reconfiguration, operating at different spatial and temporal scales in superficial and deep cortical layers.

## Methods

### Animals

All experiments were approved by the Institutional Animal Care and Use Committee (IACUC) at the Hebrew University of Jerusalem, Israel. A total of 14 adult male mice (2-6 months old) were used. Seven mice were triple-transgenic Rasgrf2-2A-dCre;CamK2a-tTA;TITL-GCaMP6f, expressing GCaMP6f in excitatory neocortical layer 2/3 neurons^40,71^ and seven were triple-transgenic Rbp4-dCre;CamK2a-tTA;TITL-GCaMP6f11, expressing GCaMP6f in excitatory neocortical layer 5 neurons (including both layer 5A and 5B neurons; Fig. 1b). These mice are the same mice used in Pollak et al. 2025^42^ but imaged on different days during expert performance (and not during learning).

To generate L2/3 triple-transgenic animals, CamK2a-tTa and TITL-GCaMP6f double transgenic mice were crossed with a Rasgrf2-2A-dCre line. Individual lines are available from The Jackson Laboratory (JAX# 016198, JAX#024103, JAX#022864). The Rasgrf2-2A-dCre;CamK2a-tTA;TITL-GCaMP6f line contain an inducible system in which the destabilized Cre (dCre), expressed under the Rasgrf2-2A promoter, can be stabilized by trimethoprim (TMP) to become functional. TMP (Sigma; T7883) was dissolved in Dimethyl sulfoxide (DMSO; Sigma; 34869) at 100 mg/ml and freshly prepared for each experiment. For induction, mice were received a single intraperitoneal injection (150 µg TMP/g body weight; 29 g needle) 3–5 days post-surgery, diluted in 0.9% saline.

To generate L5 triple-transgenic animals, the Rbp4-dCre line was crossed with the CamK2a-tTa;TITL-GCaMP6f double-transgenic line. In this line, transgene expression is suppressed by doxycycline. Because mice were not treated with doxycycline, they displayed strong expression throughout the experiment. Both lines showed layer-specific expression homogeneously across the cortex with minimal leakage into other layers, as previously reported^71,72^ (Fig. 1b).

### Surgery

We used an intact skull preparation for chronic wide-field calcium imaging of neocortical activity^40,44,73^. Mice were anesthetized with 2% isoflurane, and body temperature was maintained at 37 °C. Local analgesia (Lidocaine 1% B. Braun) was applied, the skull was exposed and cleaned, and overlying muscles were removed to access the dorsal surface of the left hemisphere (∼6 × 8 mm; ∼3 mm anterior to bregma to ∼1 mm posterior to lambda, midline to ≥ 5 mm laterally). A wall around the hemisphere was built using adhesive UV-cured material (G-Premio BOND; GC) and dental cement “worms” (Charisma; kulzer). Translucent dental cement (Tetric EvoFlow T; Ivoclar) was then applied evenly over the imaging field. Finally, a metal post for head fixation was glued to the posterior right hemisphere. This minimally invasive preparation enabled high-quality, chronic imaging of neuronal population dynamics with a high success rate.

### Texture discrimination task

Mice were trained on a whisker-based go/no-go discrimination task (Fig. 1a) using a data acquisition interface (USB-6001; National Instruments) and custom LabVIEW software^40,43,74^. Each trial began with an auditory cue (stimulus cue; 2 beeps at 2 kHz, 100 ms duration, 50 ms interval), signaling the approach of one of two sandpapers (grit size P100: rough; P1200: smooth; 3M) to the whiskers, as ‘go’ or ‘no-go’ textures (Fig. 1a). Sandpaper was mounted on panels attached to a stepper motor (X-LMS100A; Zaber), which was fixed to a motorized linear stage (T-LSM100A; Zaber) that moved textures in and out of reach. Texture contacted the whiskers for 2 s, then retracted, after which a second auditory cue (response cue; 4 beeps at 4 kHz, 50 ms duration, 25 ms interval) signaled the start of a 2 s response period. The stimulus and response cues were identical for both textures. A water reward (∼3 µL) was delivered when mice licked in response to the go texture, but only after the response cue (‘Hit’), and for the first correct lick during the response period (Fig. 1d; lick was detected using a piezo sensor). Punishment with white noise followed licking to the no-go texture (‘False Alarms’, FA). Licking before the response cue was permitted but had no consequence. No reward or punishment was delivered when mice withheld licking for the no-go (‘Correct Rejection’, CR) or go (‘Miss’) textures. The lick detector remained in a fixed, accessible position throughout the trial. Licking before the response cue was allowed and did not lead to punishment or early reward. Training and imaging were performed in complete darkness, except for blue or purple light confined to the brain that did not scatter beyond the imaging preparation. Mice could perform the task equally well with the LEDs turned off.

### Training and performance

Five mice per layer were trained to lick for the P100 texture (mice #1-4 and #7), and two mice were trained to lick for the P1200 texture (L2/3: mice #4 and #5, L5: mice #4 and #6). Mice were first handled and accustomed to head fixation before starting a water**-**restriction protocol. In this protocol, mice were deprived of water for 5 consecutive days and followed by 2 days ad libitum, and given access to water only by licking for the go texture. Mice were allowed to drink until they stopped licking and were supplemented if intake fell below 1 ml or if body weight dropped by more than 10% on consecutive days.

Before imaging, mice were conditioned to lick for a reward after the go texture, presented in a trial structure similar to the task itself. Imaging began only mice reliably licked following the response cue (typically after the 200–400 trials on the first day). On the first imaging day, mice were presented with the ‘go’ texture and after 50 trials the ‘no-go’ texture was gradually introduced (starting at 10% probability and increasing by 10% approximately every 50 trials^40^) until reaching 50% by the end of the day. After ∼100 trials, we increased the no-go probability to 80% and required three consecutive CR trials before returning to 50% probability^75^. This procedure was repeated until mice improved performance, specifically withholding licking for the no-go texture. For mice that continued to lick in response for both textures, the wrong response was repeated until a correct response was made. Training continued until animals achieved stable expert performance (d′ > 1.5) which was typically after 5 days^42^. Next, we continued imaging these mice for an additional 5-8 as they maintained high and stable performance.

### Wide-field calcium imaging

The wide-field imaging setup is consisted of a sensitive CMOS camera (Hamamatsu Orca Flash 4.0 v3; 512 x 512 pixels; 20 Hz) mounted on a dual-objective system^39,44,76,77^. Two objectives (Navitar; D-5095, D-2595) were coupled through a dichroic filter cube (510 nm; AH; Thorlabs). Blue LED light (Thorlabs; M470L3) was collimated and directed onto the preparation. Green fluorescence emitted from the preparation passes through both objectives before reaching the camera (20 Hz frame rate). An additional 405 nm light (M405L4; Thorlabs) was interleaved as a control for non-calcium dependent signals, with the normalized 405 signal subtracted from the normalized 473 signal (driven by Teensy 4.0^43,45,78^). The interleaved protocol produced a 20 Hz frame rate, yielding a 10 Hz corrected signal (see Data analysis).

### Body movement monitoring and analysis

A body camera (30 Hz frame rate; The Imaging Source; DMK 33UX273) was used to detect mild mouse movements (Fig. 1e; Supplementary Fig. 1b). For each imaging day, we manually outlined the forelimbs and back (one region of interest each), which provided reliable measures of body motion^40,43–45,76^. Body movement was calculated as 1 minus frame-to-frame correlation within these regions as a function of time for each trial, and values were averaged across both regions to generate a single ‘body movement’ vector. Forelimb and back movements during task performance were generally highly correlated with excessive whisking, which we validated using a second camera positioned below the mouse to track whisker kinematics^43^. For the linear model analysis, we manually outlined the whisk movement then calculated as 1 minus frame-to-frame correlation within these regions as a function of time for each trial to generate a single vector.

### Data analysis

Data analysis was performed using MATLAB (Mathworks). All mice were continuously imaged for 5-8 consecutive days post learning while maintaining expert performance (Fig. 1d; Supplementary Fig. 1a). Wide-field fluorescence images were downsampled to 256 × 256 pixels, and pixels outside the imaging area were discarded, resulting in a spatial resolution of ∼40 µm/pixel, sufficient to delineate cortical borders in both L2/3 and L5 despite scattering of emitted light through tissue and skull. Using an interleaved imaging protocol, signals at 470 nm and 405 nm were first extracted. For each trace, pixel and trial signals were expressed as ΔF/F, normalized to a baseline calculated from several frames preceding the stimulus cue (frame 0 division). Next, the 405 nm-normalized traces were subtracted from the 473 nm-normalized trace^43^. This correction produced minor modulation, primarily reducing post-activation dips; results were largely similar to those obtained using only the 473 nm illumination^40,43,44^.

To define regions of interests (ROIs), activation maps within each mouse were aligned across days using a semi-automatic protocol based on blood vessel patterns: 5-7 corresponding points were selected (cpselect; MATLAB) and used for transformation (cp2tform; MATLAB). Aligned maps were then registered onto the Allen Mouse Brain Atlas (©2004 Allen Institute for Brain Science, available at: http://mouse.brainmap.org/) using skull coordinates and functional patches^40,79^. Within atlas boundaries, 25 areas of interest were defined, with manual adjustments to fit each mouse’s functional activity^40,43^. Motor cortex areas were defined based on stereotaxic coordinates and functional patches (see below). Thus, ROIs were comparable both within and across mice. Although L5 top 2D view may appear slightly condensed in lateral areas, no correction was applied, as L2/3 and L5 did not seem to be substantially different^42^. The 25 areas were grouped into five categories: Motor (orange), Somatosensory (purple), Association (green), Visual (brown) and Auditory (blue) areas (Fig. 1b).

- **Motor areas:** whisker-related primary motor cortex (MOp; 1.5 anterior and 1 mm lateral from bregma, corresponding to the whisker evoked activation patch in MOp from the mapping session), anterior lateral motor cortex (ALM; 2.5 anterior and 1.5 mm lateral from bregma) and secondary motor cortex (MOs); 1.5 anterior and 0.5 mm lateral from bregma corresponding.
- **somatosensory areas:** Primary trunk (SSp-tr), Secondary sensory (SSs), Barrel cortex (SSp-bfd), Primary hindlimb (SSp-ll), Primary forelimb (SSp-ul), Primary nose (SSp-n), Primary mouth (SSp-m).
- **Association cortex:** Posterior medial (VISpm), Anterior medial (VISam), Anterior lateral (VISal), Rostro lateral (VISrl), Anterior (VISa), Restrosplenial angular (RSPagl), Restrosplenial dorsal (RSPd).
- **Visual areas:** Post rhinal (VISpor), Posterior lateral (VISpl), Lateral intermediate (VISli), Lateral medial (VISl), Primary visual (VISp).
- **Auditory areas:** Temporal association (TEa), Auditory dorsal (AUDd), Auditory primary (AUDp).

### Quantification of neural activity

The dataset comprises cortex-wide activation maps over the time course of a trial (7 seconds), recorded repeatedly across days and hundreds of trials, from seven L2/3 mice and seven L5 mice. This three-dimensional dataset (brain x trial x time^40^) can be presented as spatial activity maps (ΔF/F) averaged over trials/days and time (e.g., Fig. 2a, 4a), presented as a time course for a specific ROI and averaged over trials (e.g. Fig. 3c) or presented as a 2D map of time (seconds) versus trials for a given ROI (e.g. Fig. 5b).

### Calculation of performance curves

Trials were binned (non-overlapping bins of 30 trials) across days, and performance was calculated for each bin using d′ (d′ = Z(Hit/(Hit + Miss))−Z(FA/(FA + CR)), where Z denotes the inverse of the cumulative distribution function (Fig. 1d and Supplementary Fig. 1a).

### Quantification of Representational Drift

To quantify representational drift, population activity vectors across cortical areas were compared across days using correlation-based metrics (i.e., a vector of 25 values for each of the 5 days, representing the ΔF/F for a specific time frame/epoch). This can be done for a specific time epoch, for example within the Stimulus period (e.g. Fig 2) or for each time frame separately (e.g., Fig. 3). These analyses were done for each mouse and layer separately.

We quantified representational drift at the population level using several metrics:

- **Cross-day correlation relative to day 1:** We quantified Pearson’s correlation (R²) between the response vector on day 1 and the response vector on each subsequent day (Fig. 2c). Lower R² values indicate reduced similarity between response vectors across days^46^.
- **Rate of change in R² across days:** Using the R² values computed between day 1 and each subsequent day (yielding a vector of five R² values), we quantified the slope of the best linear fit (Fig. 2d). More negative slope values indicate a stronger degree of representational drift.
- **Correlation angle across days:** As an additional metric of representational drift, we calculated the angle between the linear fit relating day 1 to each subsequent day and the line representing a perfect correspondence (diagonal line; Fig. 1e). An increasingly negative slope of the linear fit across days indicates a progressive divergence in representations, consistent with representational drift.
- **Inter-day correlation matrices**: To assess representational stability across days, we calculated Pearson’s correlation coefficients (r) between response vectors from all possible pairs of recording days, yielding a symmetric 5 × 5 correlation matrix (Fig. 2f). Reduced correlation values with increasing distance from the diagonal indicate progressive dissimilarity between temporally distant days, consistent with representational drift. To further quantify this effect, correlation values were plotted as a function of inter-day interval (Fig. 2g), and the slope of the corresponding linear fit was extracted as a measure of drift magnitude.
- **Temporal drift measures**: The slope extracted from the inter-day correlation analysis was calculated independently for each time frame, yielding a temporal profile of drift dynamics across the trial (Fig. 3a,b). More negative slope values indicate a stronger temporal expression of representational drift at a given time point.
- **Drift within individual cortical areas:** To characterize long-term response changes within a given cortical area, we calculated the Pearson correlation coefficient (r) between response magnitude (averaged across the Stimulus period) and recording day across 30 trial bins for each cortical area (Fig. 4c). This analysis was further extended to individual time frames, yielding a temporal profile of correlation values throughout the trial structure (Fig. 5a,b). Positive r values indicate progressive increases in activity across days, whereas negative r values reflect long-term decreases in response amplitude.

### Dimensionality reduction based on laminar profiles

t-distributed stochastic neighbor embedding (t-SNE) was applied to the temporal profiles of r values across 25 cortical areas within each layer (90 × 25 matrix; cortical areas × time frames; Supplementary Fig. 6a). This analysis was performed for Hit trials, with the corresponding matrices shown in Supplementary Fig. 6a. Equivalent matrices were generated for CR trials (Supplementary Fig. 6b). The resulting t-SNE embedding provides a low-dimensional representation in which each point corresponds to a specific time frame within a given cortical area. Spatial proximity between points reflects a high degree of similarity between their corresponding high-dimensional activity patterns. We next applied k-means clustering to the embedded data, with elbow analysis indicating an optimal solution of three clusters (Fig. 6a). The cluster assignment for each cortical area and time frame is shown in Fig. 6c, and the corresponding mean r values, averaged within each cluster, are presented for each layer.

### Linear multivariate model

To assess the contribution of behavioral and task variables to neural activity, we implemented a multivariate linear model with ridge regularization using behavioral and task related predictors.

- **Trial (#):** continuous variable of the trial number.
- **Performance (d’):** continuous variable, running d-prime (30-trial bins)
- **Movement (%):** continuous variable of back and forelimb movement which was extracted from the behavioral camera (see Body movement analysis above).
- **Lick:** continuous variable of jaw movement which was extracted from the behavioral body camera.
- **Whisk:** continuous variable following whisker movement which was extracted from the behavioral whisker camera.
- **Success:** binary variable (1 = hit/correct rejection, 0 = miss/false alarm).
- **Choice:** categorical variable for the choice outcome of the mouse in each trial, converted to dummy variables (“Hit”; “Miss”; “CR”; “FA”). For example, a dummy variable for Hit will accept 1 for a Hit trial and 0 for all other choices.
- **Stimulus:** binary variable for texture type (0 = smooth/P1200, 1 = rough/P100).
- **History:** categorical variable divided to previous choice of the preceding trial, converted to dummy variables. Previous choice, a dummy variable for Hit will accept 1 if the previous trial was a Hit trial and 0 if the previous trial was any other choice.

Models were fit separately for each cortical area, time point, and animal. Regression coefficient (β) was used to quantify the strength and direction of the association between each predictor and neuronal activity, with non-zero β-values indicating a significant contribution of that predictor (Fig. 7a,b and Supplementary Fig. 7c,e).

### Variance analysis

To isolate the unique contribution of individual predictors, we computed unique explained variance (unique R²) by comparing full models to reduced models in which specific predictors were temporally shuffled. This approach allowed us to distinguish shared from independent contributions of behavioral and task variables to neural dynamics (Fig. 7c,d and Supplementary Fig. 7b,d,f).

### Statistical analysis

Statistical analyses were performed using MATLAB. Non-parametric tests, including Two-Way ANOVA, signed-rank and rank-sum tests, were used for within– and between-group comparisons, respectively. Multiple comparisons were corrected using false discovery rate (FDR) or Bonferroni procedures where appropriate. Data are reported as mean ± s.e.m. unless otherwise stated.

## Author contributions

Y.E.P and A.G conceptualized the study. Y.E.P performed the experiments and analyzed the data. Y.E.P, R.S., and A.G wrote the manuscript, and all authors revised, read and approved the final version of the manuscript.

## Data availability

Data will be made publicly available upon publication of the paper.

## Code availability

Code will be made publicly available upon publication of the paper.

## Supplementary materials

Figures S1 to S7

## Acknowledgements

We would like to thank Malak Abumadi and Dr. Odeya Marmor for their help with genotyping and mouse maintenance, Fritjof Helmchen and Philipp Bethge for the help with the transgenic mice lines. This work is funded by the Einstein Foundation Research Grant (A-2021-644; A.G and M.L) and the European Union (ERC Starting Grant, MESO-AG, 101040378).

