## Supplementary figures for "Cortex-wide representational drift of different layers"

**Supplementary material**

Supplementary Figures 1-7.

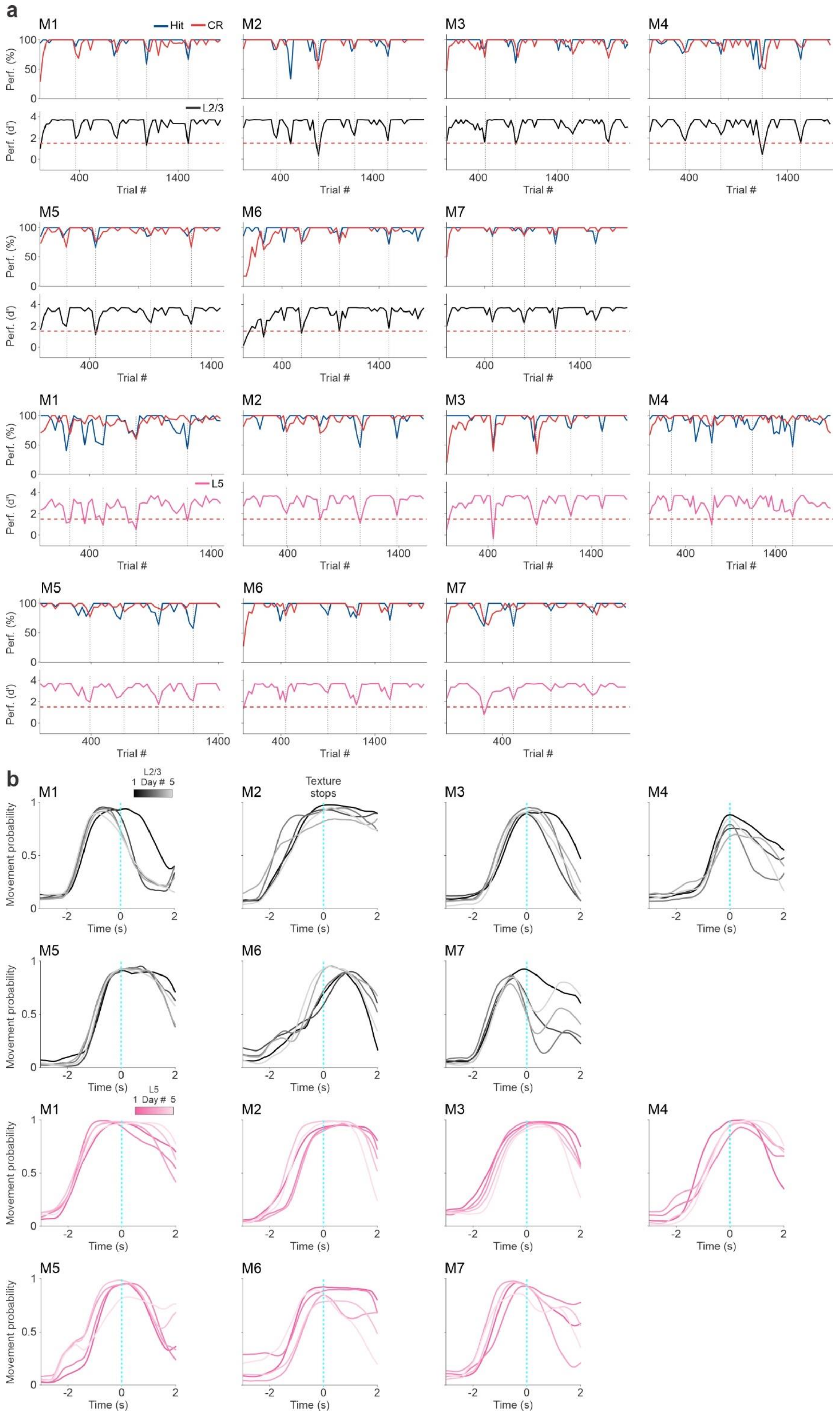

a
M1
M2
M3
M4
M5
M6
M7
Hit
CR
L2/3
L5
Perf. (%)
Perf. (d')
Trial #
b
L2/3
1 Day # 5
L5
Texture stops
Movement probability
Time (s)

**Supplementary Fig. 1 Performance and movement are similar across days in individual mice. a** *Top:* Percentage (%) of Hit (blue) and CR trials (red) as a function of trials (30 trial bins) for each mouse separately. *Bottom:* Performance (d') as a function of trial number for each mouse separately: L2/3 (black; n = 7) and L5 (pink; n = 7). Red dashed horizontal line indicates threshold for learning (d' = 1.5), gray dashed vertical lines indicate the end of every recording day. **b** Movement probability along the temporal trial structure for each day (color codded chronologically from dark to light shades) for each mouse in L2/3 (black; n = 7) and L5 (pink; n = 7). Vertical cyan dashed lines depict the texture stop.

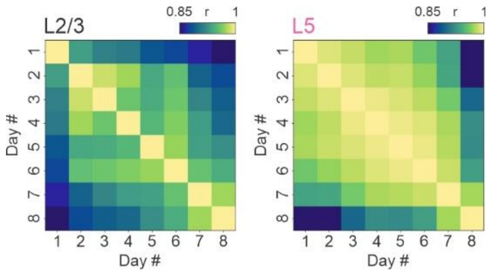


**Supplementary Fig. 2 Inter-day correlation matrix for 8 consecutive days.** Average population vector of inter-day correlations for 25 cortical areas (as in Fig. 2f) for one example mouse in L2/3 (left) and L5 (right) across 8 consecutive recording days. Color scale bar indicates min/max of Pearson's correlation coefficient (r). In general, a similar pattern remains between 8-by-8 and 5-by-5 matrices.

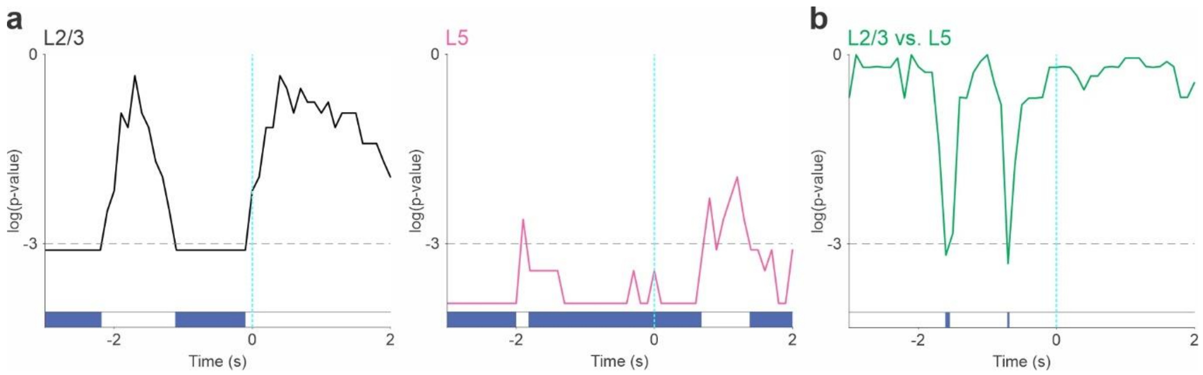


**Supplementary Fig. 3 Frame-by-frame significance test for representational drift within and across layers. a** Frame-by-frame p-value (in log scale) derived from a statistical comparison between slope (presented in Fig. 3b bottom) and 0 (signed-rank test; FDR-corrected; n = 7 mice for each layer). L2/3 (left; black) and L5 (right; pink). Significance threshold (log(p) = -3) is marked in a gray dashed line. A binary vector indicating significance (blue) is presented below. **b** A frame-by-frame statistical test between the slopes in L2/3 and L5 (green; rank-sum test; FDR-corrected; n = 7 mice for each layer; corresponding to the plot in Figure 3b bottom).

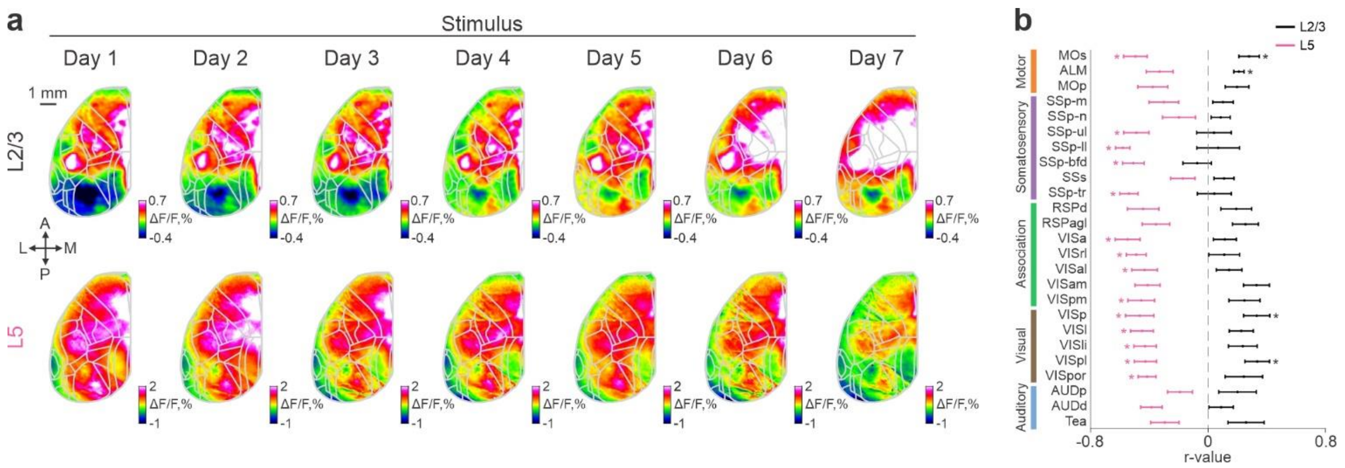


**Supplementary Fig. 4 Additional analysis for cortex-wide activity across time during early stimulation.** Example activation maps from one mouse of L2/3 (top) and L5 (bottom) averaged during early stimulation (Stimulus; -0.4 to 0 seconds relative to texture stop) and across 7 consecutive days (left to right; Compared to 5 days in Fig. 4a). Color scale bar indicates min/max of percent ΔF/F. Overlay of areas in gray for all maps. Scale bar is 1 mm. **b** Pearson's correlation coefficient (r; ΔF/F as a function of trial bins; As in Figure 5) across 25 cortical areas during Stimulus averaged across L2/3 (black) and L5 (pink) mice. Error bars are ± s.e.m. across mice (n = 7). *$p$ < 0.05, signed-rank test, FDR-corrected.

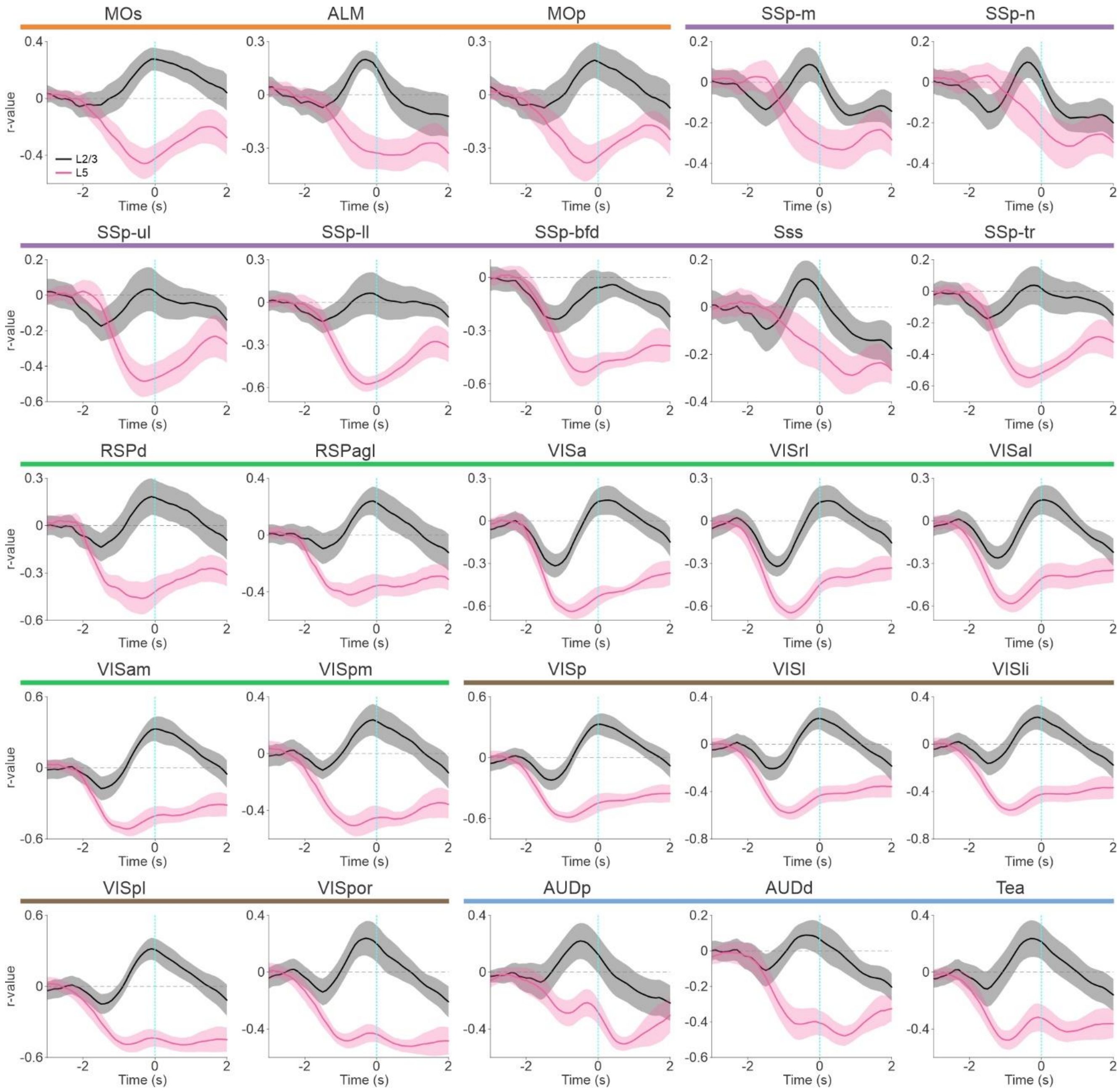


**Supplementary Fig. 5 Laminar representational drift temporal relation to trial number across 25 cortical areas.** Pearson's correlation coefficient (r) curve averaged across L2/3 (black) and L5 (pink) mice as in Figure 5b (bottom) but for all 25 cortical areas. Error bars are ± s.e.m. across mice (n = 7). Vertical cyan dashed lines depict the texture stop. Regions are grouped into five divisions: Auditory (blue), Visual (brown), Association (green), Somatosensory (purple), and Motor (orange).

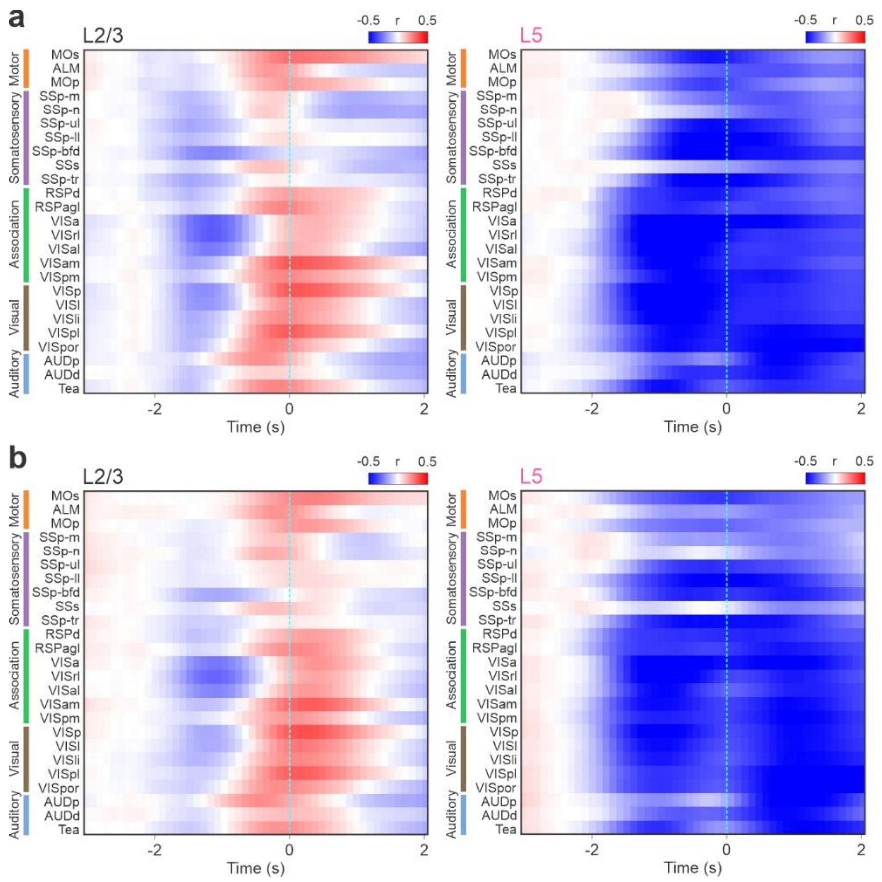


**Supplementary Fig. 6 Dimensionality reduction extended data. a** The input matrices used for the t-SNE dimensionality reduction in Figure 6a. The input is a 2D heat map of 25 cortical areas by time frames (n = 90) where each element is the Pearson's correlation coefficient (r; as in Figure 5b and Supplemental Figure 5; Hit trials) averaged across L2/3 (left) and L5 mice (right). Vertical cyan dashed lines depict the texture stop. Color scale bar indicates min/max of r **b** Similar heat maps but for CR trials, showing similar dynamics.

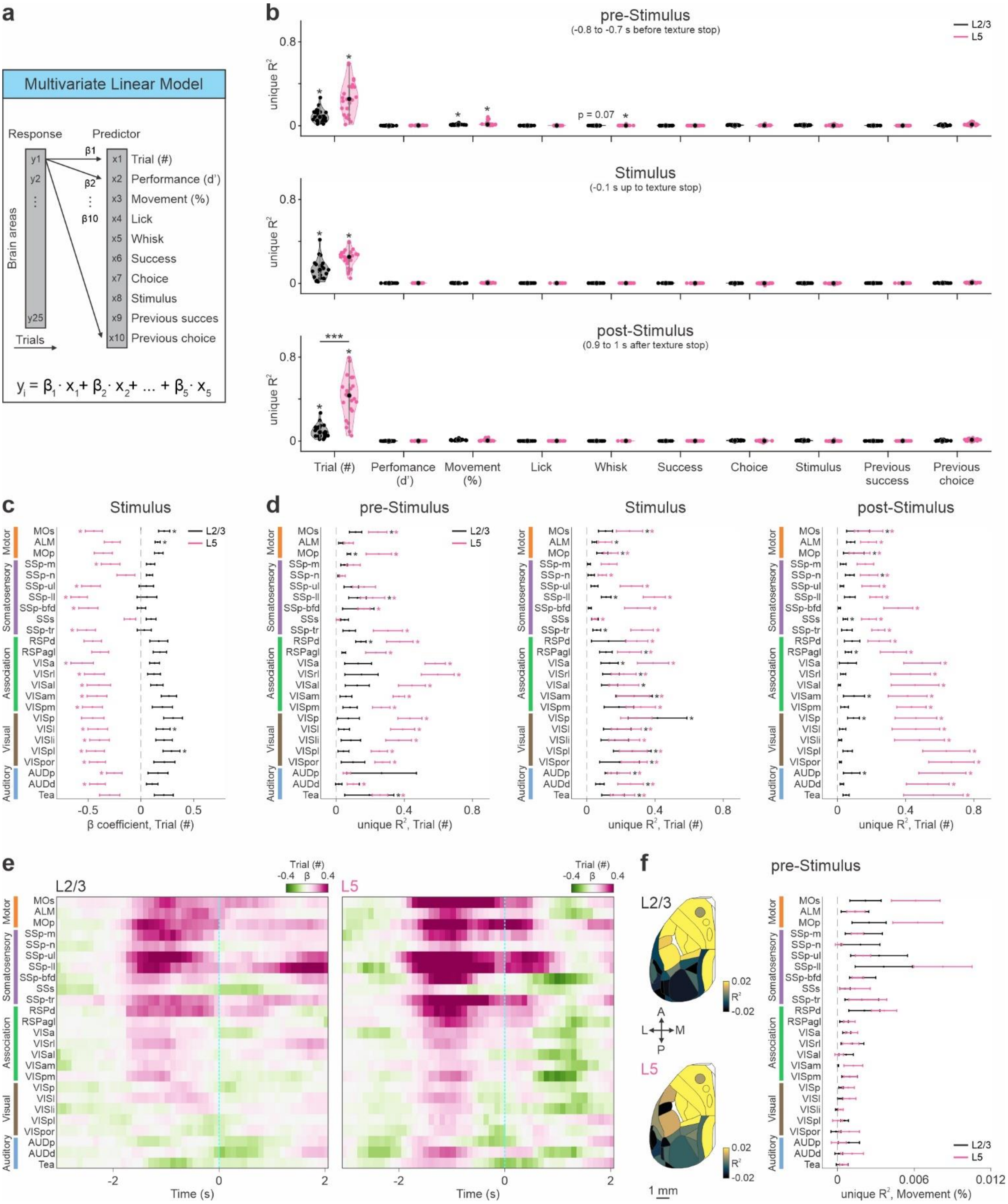
a
Multivariate Linear Model
Response
Predictor
Brain areas
y1
y2
y25
Trials
β1
β2
β10
x1 Trial (#)
x2 Performance (d')
x3 Movement (%)
x4 Lick
x5 Whisk
x6 Success
x7 Choice
x8 Stimulus
x9 Previous succes
x10 Previous choice
$y_i = \beta_1 \cdot x_1 + \beta_2 \cdot x_2 + \dots + \beta_5 \cdot x_5$
b
pre-Stimulus
(-0.8 to -0.7 s before texture stop)
Stimulus
(-0.1 s up to texture stop)
post-Stimulus
(0.9 to 1 s after texture stop)
unique R²
L2/3
L5
p = 0.07
***
Trial (#)
Perfomance (d')
Movement (%)
Lick
Whisk
Success
Choice
Stimulus
Previous success
Previous choice
c
Stimulus
β coefficient, Trial (#)
d
pre-Stimulus
Stimulus
post-Stimulus
unique R², Trial (#)
Motor
Somatosensory
Association
Visual
Auditory
MOs
ALM
MOp
SSp-m
SSp-n
SSp-ul
SSp-ll
SSp-bfd
SSs
SSp-tr
RSPd
RSPagl
VISa
VISrl
VISal
VISam
VISpm
VISp
VISl
VISli
VISpl
VISpor
AUDp
AUDd
Tea
e
L2/3
L5
Trial (#)
β
Time (s)
f
pre-Stimulus
A
L
M
P
R²
1 mm
unique R², Movement (%)

**Supplementary Fig. 7 Additional analysis for the multivariate linear regression. a** Schematic illustration of a multivariate linear model that predicts neuronal responses in each brain area (as a function of time) from a set of behavioral and task related predictors **b** Violin plot of unique $R^2$ for individual predictors averaged across three selected temporal periods: pre-Stimulus, Stimulus and post-Stimulus (from top to bottom) and mice of L2/3 (black; n = 7) and L5 (pink; n = 7). Dots depict 25 different brain areas median across mice (n = 7). *$p < 0.05$, ***$p < 0.0005$ sign-rank test within layers. rank-sum test across layers, Bonferroni-corrected. **c** Coefficient value (β) for the trial number predictor averaged across all 25 cortical areas for L2/3 (black) and L5 (pink) mice during Stimulus corresponding to Figure 7b. Error bars are ± s.e.m. across mice (n = 7). *$p < 0.05$, signed-rank test, FDR-corrected. **d** Unique $R^2$ for the trial number predictor averaged within the three selected temporal periods: pre-Stimulus, Stimulus and post-Stimulus (from left to right) for L2/3 (black) and L5 (pink) mice. Error bars are ± s.e.m. across mice (n = 7). *$p < 0.05$, signed-rank test, FDR-corrected. **e** Coefficient values (β) averaged for the movement probability predictor (move. (%)) as cortical activity maps of 25 cortical areas by time frames (n = 90) for L2/3 (left) and L5 (right) mice. Color scale bar indicates min/max of Coefficient value (β). Vertical cyan dashed lines depict the texture stop. **f** *Left:* Cortical maps of unique $R^2$ for the movement probability predictor averaged across pre-Stimulus and L2/3 and L5 mice. Color scale bar indicates min/max of unique $R^2$. *Right:* Unique $R^2$ for movement probability predictor averaged across pre-Stimulus in L2/3 (black) and L5 (pink) mice. Error bars are ± s.e.m. across mice (n = 7).
